# Integrating flowering and stress responses in Arabidopsis through KH-domain genes

**DOI:** 10.1101/2025.01.09.631668

**Authors:** Encarnación Rodríguez-Cazorla, Juan-José Ripoll, Héctor Candela, Almudena Aranda-Martínez, Ernesto Zavala-González, Antonio Martínez-Laborda, Antonio Vera

## Abstract

Plant reproductive success largely relies on flowering under favorable conditions. However, stress factors have forced plants to acquire adaptive strategies to coordinate floral timing and stress responses through key genetic elements. RNA-binding proteins with K-homology (KH) domains are emerging as versatile regulators of an increasing number of plant developmental processes, including flowering and stress acclimation. In *Arabidopsis thaliana*, *FLK* and *HOS5* encode multifaceted KH-domain proteins associated with transcription and cotranscriptional operations. *FLK* facilitates floral transition by repressing *FLC*, the central flowering inhibitor, while both KH-genes have been involved in abiotic stress and defense against pathogens. Our genetic and molecular data identify HOS5 as a novel flowering regulator that, together with FLK, represses *FLC*. Our transcriptomics results reveal that, in addition, *FLK* and *HOS5* cooperatively repress numerous stress-responsive genes. Consistent with this, *flk hos5* double mutant plants exhibit elevated levels of stress-induced gene activities and enhanced resistance to abiotic stress and pathogenic fungi. The coordinated repression of *FLC* and stress-induced genes by FLK and HOS5 suggests that these KH proteins are part of a cotranscriptional regulatory hub key for orchestrating flowering time and environmental adaptation responses.

## Introduction

To initiate flowering under optimal conditions, plants have evolved a sophisticated network of regulatory pathways that sense environmental and endogenous cues (photoperiod, temperature, developmental phase), that finally converge on floral integrators, triggering the formation of flowers (Kinoshita and Richter, 2020; Freytes *et al*., 2021; Quiroz *et al*., 2021) In the reference plant *Arabidopsis thaliana* (Arabidopsis hereafter), the autonomous pathway (AP) genes promote flowering independently of daylength by repressing the central flowering inhibitor *FLOWERING LOCUS C* (*FLC*) (Wu et al., 2020). *FLC* encodes a MADS-domain transcription factor that prevents precocious flowering by directly repressing floral integrators such as *FLOWERING LOCUS T* (*FT*, the florigen) and *SUPPRESSOR OF OVEREXPRESSION OF CONSTANS1* (*SOC1*) (Michaels & Amasino, 1999; Li et al., 2008; Jang et al., 2009).

In addition to daily and seasonal ambient fluctuations, plants are often exposed to biotic (pathogenic microorganisms), or abiotic stress conditions (including drought, salinity, or extreme temperatures) which are further exacerbated by global climate change, compromising survival and reproduction (Fichman & Mittler, 2020). To neutralize these effects, plants have developed complex defense mechanisms that balance stress-tolerance and growth, ultimately affecting yield (Park *et al*., 2016; Zhang *et al*., 2020a). One significant response to stress is the modification of flowering time through an intricated genetic and molecular network connecting both processes (Kazan & Lyons, 2016). For example, salinity usually delays flowering whereas drought can accelerate it. Indeed, stress and flowering pathways ultimately converge on common endogenous floral regulators (Kazan & Lyons, 2016). For instance, short periods of cold stress delay flowering by activating *FLC* (Jung et al., 2013) and the AP genes *FPA* and *FLOWERING LOCUS D* (*FLD*) negatively regulate resistance against the bacterial pathogen *Pseudomonas syringae* (Lyons et al., 2013; Singh et al., 2013). Therefore, dissecting the genetic mechanisms underlying the crosstalk between flowering and stress pathways is crucial for understanding the adaptation of floral timing to challenging environmental conditions.

As sessile organisms, plants must respond to environmental variations through rapid changes in gene regulation. Transcriptional reprogramming and associated cotranscriptional pre-mRNA processing, including 5’ capping, splicing, and 3’ cleavage/polyadenylation, are major determinants of gene expression, and also generate multiple isoforms that increase developmental flexibility and adaptive responses (Ambrosone et al., 2012; Marquardt et al., 2023). RNA-binding proteins are crucial to accomplish these functions (Ambrosone et al., 2012; Bentley, 2014; Rodríguez-Cazorla et al., 2015; Rodríguez-Cazorla et al., 2018; Marquardt et al., 2023; Shine et al., 2024). *FLOWERING LOCUS K* (*FLK*) encodes an RNA-binding protein with K-homology (KH) domains that, as an AP member, promotes flowering via *FLC* suppression (Lim et al., 2004; Mockler et al., 2004). The KH domain, originally identified in the human heterogeneous nuclear RNP K (hnRNPK; Siomi et al., 1993), is an ancient motif important for binding to single-stranded DNA/RNA, and provides a structural basis for protein-protein interactions (Makeyev & Liebhaber, 2002; Nicastro et al., 2015). Thus, proteins with KH domains are involved in all levels of gene regulation and disruption of KH-domain genes are linked to severe human disorders (Lewis et al., 2000; Makeyev & Liebhaber, 2002; Hasan & Brady, 2024). The structure of FLK, harboring three-KH motifs, closely resembles that of metazoans Poly(rC)-Binding Proteins (PCBP), a functionally versatile family of proteins that includes hnRNPK (Lim et al., 2004; Zhao et al., 2022). Interestingly, a recent study identified FLK as an N6-methyladenosine (m^6^A) reader that represses *FLC* by reducing mRNA stability and splicing (Amara et al., 2023). Additionally, previous evidence also suggested a transcriptional role for *FLK* via chromatin modulation (Veley & Michaels, 2008).

During flower morphogenesis, *FLK* acts in concert with two other KH genes, *PEPPER* (*PEP*) and *HUA ENHANCER4* (*HEN4*), to secure the correct expression of the floral master regulator *AGAMOUS* (*AG*), a MADS-box encoding gene similar to *FLC* (Cheng et al., 2003; Rodríguez-Cazorla et al., 2015). FLK, PEP and HEN4 interact at the protein level, suggesting that they participate in the same complexes to regulate their targets cotranscriptionally (Rodríguez-Cazorla et al., 2015). However, *PEP* and *HEN4* promote *FLC* expression, thus antagonizing *FLK* during flowering regulation (Ripoll et al., 2009; Ortuño-Miquel et al., 2019). This suggests that the function and/or composition of common ribonucleoprotein assemblies is dynamic and complex, and most probably involving yet unknown additional partners.

In addition to flowering and floral morphogenesis, *FLK* has been recently linked to pathogen defense and salicylic acid (SA) homeostasis (Fabian et al., 2023), making this gene an appealing candidate to coordinate stress responses and adaptation of reproductive development. However, the mechanisms by which *FLK* links both operations remain unclear. The identification of additional *FLK* interacting factors might reveal additional insights into our understanding about these processes. The gene *HIGH OSMOTIC STRESS GENE EXPRESSION 5* (*HOS5*), also known as *SHINY1* (*SHI1*), *REGULATOR OF CBF GENE EXPRESSION3* (*RCF3*), or *ENHANCED STRESS RESPONSE1* (*ESR1*), encodes another KH-domain protein involved in abiotic stress and pathogen resistance (Xiong et al., 1999; Chen et al., 2013; Guan et al., 2013; Jiang et al., 2013; Karlsson et al., 2015; Thatcher et al., 2015; Jeong et al., 2013). HOS5 has been proposed to regulate splicing, and repress transcription of stress inducible genes by preventing mRNA capping, and thus the transition to transcript elongation (Chen et al., 2013; Jeong et al., 2013; Jiang et al., 2013).

To better delineate the role of *FLK* in flowering adaptation and stress regulation, we have explored its relationship with *HOS5*. Strong genetic interactions evidence that both genes act in concert to repress *FLC* expression, unveiling *HOS5* as a novel flowering regulator. We also show that *FLK* and *HOS5* coordinately coregulate numerous genes involved in stress responses. In line with this, *flk hos5* double mutants show upregulation of numerous “stress genes”, elevated levels of the defense hormones jasmonic acid (JA) and SA, and higher tolerance to abiotic stressors and pathogenic fungi. Our genetic and molecular data support a model in which FLK and HOS5 directly cooperate as part of a regulatory module that integrates plant developmental outputs (flowering) and environmental (stress) adaptive responses, a view reinforced by the ability of FLK to interact physically with HOS5. We also discuss possible mechanisms by which *FLK* and *HOS5* interact to regulate mRNA expression of *FLC* and additional gene targets. This study further expands our knowledge on the underlying molecular mechanisms governing flowering and stress response coordination, and provides candidates for their exploitation in crop biotechnological strategies.

## Materials and methods

### Plant material

All strains in this work were in the Arabidopsis Columbia (Col-0) accession: *flk-2* (Mockler et al., 2004), *hos5-2* (Chen et al., 2013), *hos5-5* (SALK_013918, this work), *flc-3* (Michaels & Amasino, 1999). Information about molecular genotyping and primers used in this work can be found in Table S1.

### Standard growth conditions and flowering time measurements

Surface-sterilized seeds were germinated on half MS plates at 21°C under long-day (16h-8h) or short-day (8h-16h) regimes, as previously described (Ripoll et al., 2009; Zavala-Gonzalez et al., 2017). Fourteen-day-old seedlings were transplanted to individual pots with soil, and inspected daily for flowering (days and rosette leaves at bolting). Except otherwise indicated, 30 plants per genotype were analyzed in a single assay, and every experiment was carried out three times.

### Germination and growth under salt and methyl-viologen

Seeds were sown on medium supplemented with varying concentrations of NaCl, under long-day conditions. Germination was determined counting radicle emergence under a dissecting microscope. For methyl viologen (MV; Paraquat) treatments, seeds were plated onto media with 0.5 or 1 μM MV (Sigma-Aldrich), and seedlings with green, fully emerged cotyledons were counted. A minimum of 100 seeds per replicate were scrutinized for each genotype under analysis. Standard deviation (SD) was calculated from three independent experiments, except for growth at 50 mM NaCl (SD calculated from two replicates with 12 plants per genotype).

### Methyl-jasmonate (MeJA) root inhibition assays

Seeds were grown on vertically-oriented control plates or supplemented with 50 µM MeJA. Seven-day-old plants were photographed and primary root length was determined using Image J software. Three independent experiments were conducted with 20 plants per genotype in each assay.

### Quantitative reverse transcription PCR (qRT-PCR)

All RNA extractions were conducted at Zeitgeber Time (ZT) 3 (h). qRT-PCR procedures (5 μg of total RNA extracted from 12-day-old rosettes) were as previously reported (Rodríguez-Cazorla et al., 2020). For each experiment, three biological replicates were performed, with three technical replicates each. Splicing efficiency was determined, for each intron examined, as the level of spliced transcript normalized to the amount of unspliced transcript, and represented as the fold change over Col-0 values from three independent assays.

### RNA-Seq and bioinformatics analysis

Total RNA was extracted (Rodríguez-Cazorla et al., 2015) from pooled 12-day-old rosettes. One μg RNA per sample was used for cDNA library construction with the TruSeq Stranded mRNA LT Sample Prep Kit for Illumina® (NEB, USA). The resulting fragments were sequenced in the lllumina Hiseq 2500 platform, using 151 bp paired-end reads, at Macrogen (South Korea). Paired-end reads were first processed using Trimmomatic v. 0.39 (Bolger et al., 2014) with options ILLUMINACLIP:TruSeq3-PE.fa:2:30:10:2:keepBothReads LEADING:3 TRAILING:3 MINLEN:36. The reads were then aligned to the TAIR10 version of the *Arabidopsis thaliana* genome sequence (https://www.arabidopsis.org/) using Hisat 2 version 2.2.1 (Kim et al., 2019), considering the strandness of the reads (with option --rna-strandness RF) and discarding all discordant read mappings (with options no-discordant and no-mixed). Transcript levels were quantified for the ARAPORT11 gene models using the cuffdiff program of the Cufflinks version 2.2.1 package (Trapnell et al., 2013), selecting fr-firststrand as the library type. HTSeq-count (version 2.0.5; Anders et al., 2015) was used to count reads aligned to introns, with the following parameters: -f bam -r pos -s reverse --nonunique all -t intron -i gene_id. The resulting counts were analyzed using the DESeq2 package (version 1.38.3; Love et al., 2014) in R (version 4.2.2). Introns with an adjusted p-value < 0.05 and an absolute log2 fold change > 1 were considered significantly differentially expressed. Three biological replicates were used for each genotype. The resulting read alignments, supplied as files in BAM format, were visualized using Integrative Genomics Viewer (IGV) software (Thorvaldsdottir et al., 2013). Identification of overrepresented GO terms was performed as implemented by the Panther classification system in The Gene Ontology website (http://geneontology.org/) using a selected set of genes (including those marked “OK” by Cufflinks) as the customized annotated reference, as previously described (Muñoz-Nortes et al., 2017). Fisher’s Exact was used as test type with the Bonferroni correction for multiple testing.

### Protein interactions

Bimolecular fluorescence complementation (BiFC) and yeast two hybrid (Y2H) assays were performed as previously described (Ripoll et al., 2015; Guan et al., 2017; Rodríguez-Cazorla et al., 2018). The corresponding constructs were generated via Gibson DNA assembly (Gibson, 2011). Empty vectors were used as negative controls.

### Quantification of plant hormones

Measurements of jasmonate (JA) and salicylic acid (SA) were carried out as in Zavala-Gonzalez et al. (2017), according to Seo et al. (2011). For every measurement, we used 100 mg of plant material (12-day-old pooled rosettes). At least 30 plants per sample were used and the experiment was carried out three times.

### Fungal inoculation

Fungal strains *Botrytis cinerea* (BC03, CECT No. 20973, IRTA Institute, Spain), and *Fusarium oxysporum* (EAN 350, CECT No. 2154) were maintained in Potato Dextrose Agar (PDA), and subcultured monthly. Conidia were obtained from 25-day-old colonies on PDA plates using 0.02% Tween 20. Resulting conidia suspensions were filtered with glass wool, counted using a haemocytometer, and adjusted to 10^5^ spores per milliliter. A drop (10 µl) of spore solution was applied on the top of each two-week-old plant grown on agar plates (long-day), and photographed 15 days after inoculation (dai). Forty plants per genotype were examined, and the experiments were repeated three times.

### Statistics

Data were subjected to analysis of variance (ANOVA) to determine significant differences among genotypes (\**P* < 0.05; \*\**P* < 0.01; \*\*\**P* < 0.001). SD was calculated in Microsoft EXCEL from aggregate data from independent experiments. For qRT-PCR experiments, relative expression was calculated according to Pfaffl et al. (2002), and statistical significance was estimated by Student’s t-test (\**P* < 0.05; \*\**P* < 0.01; \*\*\**P* < 0.001).

## Results

### *HOS5* interacts with *FLK* to repress *FLC* and promote flowering

To explore the connections between *FLK* and *HOS5*, we generated double mutants using the null T-DNA alleles *hos5-2*, *hos5-5* (Figure S1) and *flk-2* (Mockler et al., 2004). None of the single or double mutants showed any conspicuous morphological defect when compared to wild-type Col-0 plants (Figure S2). However, under long-day conditions, *hos5* mutants flowered similar or slightly later than Col-0, whereas, as previously reported (Lim et al., 2004), *flk-2* plants flowered much later (Figure 1a,b). Strikingly, flowering was dramatically delayed in *flk-2 hos5* double mutants with respect to *flk-2* (Figure 1a,b). On average, *flk-2* plants flowered after 30 days and 25 leaves, whereas *flk-2 hos5* double mutants required more than 45 days and 40 leaves to bolt (Figure 1b), revealing a synergistic interaction between *FLK* and *HOS5*. Under short-day conditions, *flk-2 hos5-5* plants also flowered significantly later than *flk-2* (Figure S3). This strong genetic interaction uncovers a new role for *HOS5*, in concert with *FLK*, in flowering time regulation.

**Figure 1.**
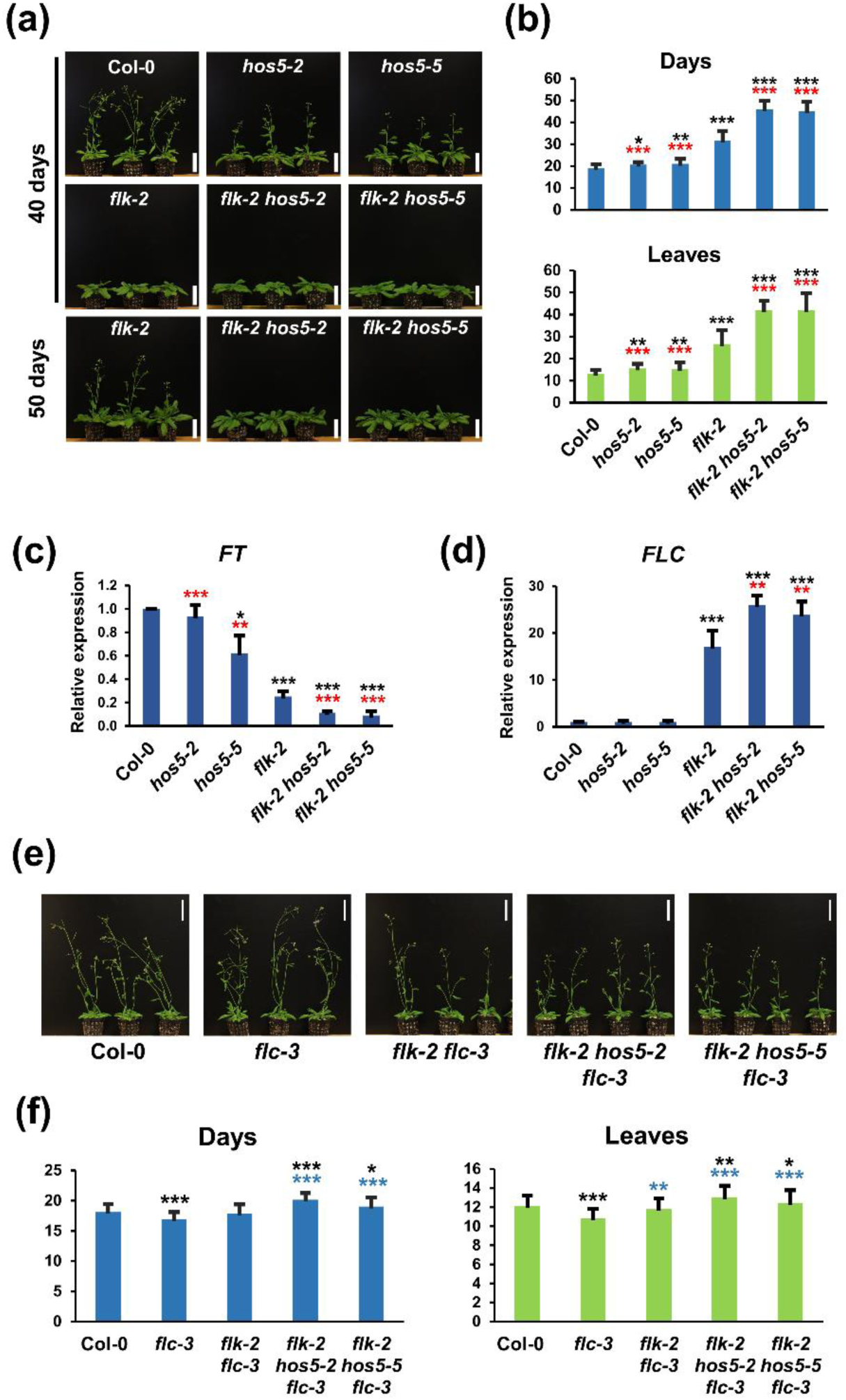
*FLK* and *HOS5* interact to promote flowering via *FLC* repression. (a) Representative 40- and 50-day-old Col-0 (wild type), and mutant plants. (b) Flowering time of Col-0 and mutant plants (number of days or rosette leaves at bolting). (c,d) Relative expression of *FT* (c) and *FLC* (d) monitored by qRT-PCR. Data correspond to three biological replicates with three technical replicates each. Bars represent mean ± SD (standard deviation). (e) Representative 37-day-old Col-0 and mutant plants harboring *flc-3*. (f) Flowering time at bolting for each of the corresponding backgrounds. For flowering assays (b, f), bars indicate mean ± SD from three independent experiments, with at least 30 plants per genotype each. For flowering and/or qRT-PCR, black, red and blue asterisks indicate significant differences with respect to Col-0, *flk-2* and *flc-3,* respectively (*, *P* < 0.05; **, *P* < 0.01; ***, *P* < 0.001). Scale bars, 5 cm.

In *flk*, plants flower late due to *FLC* overexpression (Lim et al., 2004). *FLC* presents four isoforms, being variant 1, by far, the most abundant (Cai et al., 2023). We monitored, by qRT-PCR, *FLC* expression as the amount of transcript corresponding to correctly spliced intron 1, common to all four isoforms. In *hos5* mutants, *FLC* mRNA levels were largely similar to those of Col-0. However, in *flk-2 hos5* plants, *FLC* abundance was significantly higher than in *flk-2*. These results closely correlate with the observed delay in flowering time (Figure 1b,d), and suggest that *FLC* misexpression is the likely cause of this phenotype. Consistent with this, the expression of integrator genes repressed by *FLC* was significantly downregulated in *flk-2* and *flk-2 hos5* mutants (Figure 1c; Figure S4). These findings were consistently observed in both combinations of *flk-2 hos5* double mutants (Figure 1; Figure S4), indicating that *hos5-2* and *hos5-5* are equivalent null alleles. We therefore adopted *hos5-5* as the reference hereafter.

To confirm the direct involvement of *FLC* in the flowering phenotypes, we introduced the *flc-3* null allele (Michaels & Amasino, 1999) into both *flk-2* and *flk-2 hos5-5*. In the resulting backgrounds, flowering delay was abolished (Figure 1e,f), providing genetic evidence that the late-flowering phenotypes of *flk-2 hos5* mutants result from *FLC* upregulation. However, a modest but significant delay in the *flk-2 hos5-5 flc-3* mutants, as compared to Col-0 and *flc-3* individuals, respectively, suggests the existence of *FLC*-independent effects (Figure 1f).

### High levels of *FLC* expression mediate the germination vigor of *flk hos5* seeds

*FLC* inhibits flowering, but positively regulates other developmental processes, such as the germination transition, making its expression a pleiotropic trait of significant adaptive relevance. *FLC* enhances germination efficiency by modulating gene activities that reduce the germination-repressive hormone abscisic acid and trigger the germination-inductive gibberellins (Chiang et al., 2009). Consistent with this, high *FLC*-expressing strains, such as *flk hos5* double mutants, often exhibit robust germination. Therefore, to further support our observations on flowering time, we tested wild-type and mutant germination under salt stress.

Germination was scored on agar medium with increasing NaCl concentrations. In control plates, all genotypes rapidly reached 100% of germination rates, and salinity correlated with lower germination percentages (Figure 2a). Under salt stress, Col-0 and *hos5-5* exhibited similar poor germination rates, whereas *flk-2* and *flk-2 hos5-5* seeds germinated more vigorously. Strikingly, at 300 mM NaCl, *flk-2 hos5-5* seeds exhibited 20% germination rate, while it was completely inhibited for the rest of the genotypes assayed (Figure 2a).

**Figure 2.**
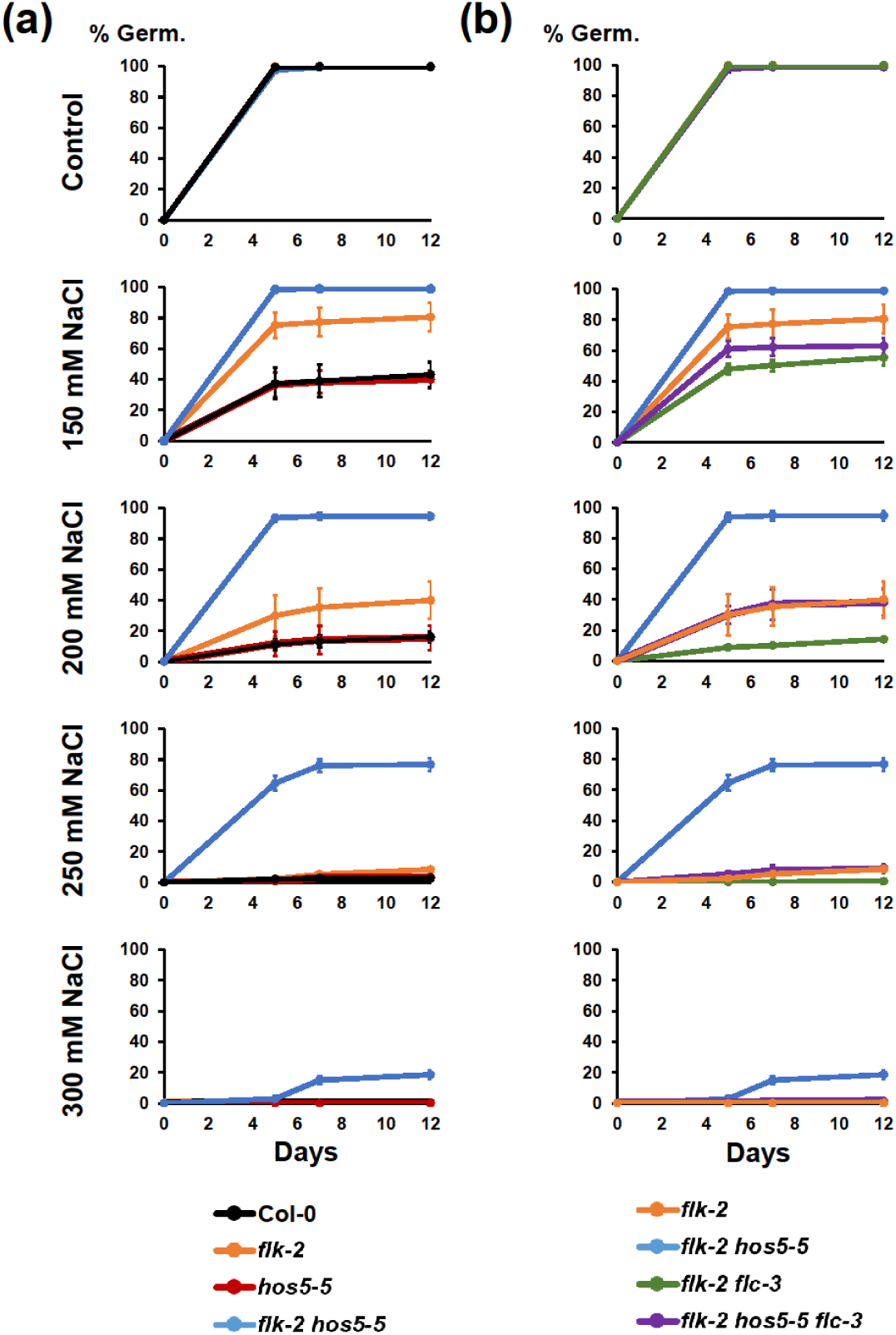
*FLC* mediates *flk hos5* germination vigor under salt stress. (a) Percentage of germination in the wild type Col-0, and *flk-2, hos5-5*, and *flk-2 hos5-5* mutant backgrounds. (b) Percentage of germination in the *flk-2 flc-3* and *flk-2 hos5-5 flc-3* mutant backgrounds. For a better comparison, and underscore the relevance of *FLC* on their elevated germination rates, the same *flk-2* and *flk-2 hos5-5* data shown in panel a) are also included in b). The appearance of visible radicles was used as a morphological marker for germination. Three independent measurements, with no less than 100 seeds, were averaged. Bars indicate mean ± SD.

Germination rates of the *flk-2* and *flk-2 hos5-5* genotypes are consistent with our hypothesis, and nicely correlated with higher *FLC* mRNA levels observed during seedling development. Therefore, to further substantiate this issue we studied *flk-2 flc-3* and *flk-2 hos5-5 flc-3* germination under salt stress. The lack of *FLC* greatly reduced the ability to germinate in saline medium. In *flk-2 flc-3*, germination rates plummeted to wild-type values (Figure 2b). Similarly, the *flk-2 hos5-5 flc-3* seed germination rates were much lower than those of *flk-2 hos5-5*. However, *flk-2 hos5-5 flc-3* triple mutants still germinated slightly better than Col-0 seeds (Figure 2b). This indicates that, although *FLC* accounts for a large part of the high germination rate for *flk-2 hos5-5* seeds, *FLC*-independent factors contribute to a minor fraction of their germination vigor, echoing our observations on flowering. These results reinforce the importance of the *FLK-HOS5* interaction for *FLC* regulation and extend its relevance to another fundamental aspect of plant reproduction: seed germination.

### Genome-wide profiling suggests that *FLK* and *HOS5* interact to limit the expression of stress-inducible genes

In addition to regulating *FLC*, *FLK* and *HOS5* participate in stress and defense responses (Xiong et al., 1999; Chen et al., 2013; Guan et al., 2013; Jiang et al., 2013; Thatcher et al., 2015; Fabian et al., 2023), suggesting that FLK-HOS5 likely constitute a regulatory hub that integrates flowering and stress response pathways. To delve into the transcriptomic landscape influenced by *FLK* and *HOS5*, we performed RNA sequencing (RNA-Seq) experiments. RNA was isolated from Col-0, *flk-2*, *hos5-5* and *flk-2 hos5-5* plants grown under long-day conditions. Our RNA-Seq analysis pipeline (false discovery rate, FDR, threshold of 5%) uncovered numerous differentially expressed genes (DEG) relative to the wild type (Table S2). We identified 762 and 433 genes less expressed in *flk-2* and *hos5-5*, respectively, including 189 common genes (Figure S5). We also found 590 significantly overexpressed genes in *flk-2*, and 815 in *hos5-5*, being 356 common to both groups (Figure S5), indicating that *FLK* and *HOS5* share common downstream genes.

Interestingly, we detected 3348 DEGs in the *flk-2 hos5-5* double mutant, nearly three times the number found in each single mutant. Among them, 1533 loci were downregulated whereas other 1815 were significantly overexpressed above wild-type levels (Figure S5; Table S2). Furthermore, the striking increase of DEGs in *flk-2 hos5-5* plants revealed a high number of genes specifically altered in the double mutant (986 down- and 1024-upregulated loci; Figure S5). All together suggests that *FLK* and *HOS5* are broad-spectrum regulatory genes and emphasizes the relevance of their interaction. Transcriptomic profiling was validated by qRT-PCR expression analyses of *FLC* and additional genes, which largely mirrored RNA-Seq abundance profiles (Figure 1, Figure S4; Figure S6; Table S2).

To further our understanding of the processes influenced by *FLK* and *HOS5*, we performed an enrichment analysis using The Gene Ontology (GO) database. Many overrepresented GO terms for biological processes were related to stress responses (Table S3). Enriched GO terms, such as ‘cellular response to hypoxia’, ‘response to cold’, ‘defense response to fungus’, ‘defense response to bacterium’, ‘response to salicylic acid’ or ‘response to oxidative stress’ were identified from upregulated genes for the three mutant backgrounds evaluated. The *hos5-5* and *flk-2 hos5-5* mutants also showed enrichment for the GO term ‘innate immune response’, whereas both *flk-2* and *flk-2 hos5-5* exhibited enrichment in ‘response to salt stress’, and JA-associated GO terms (Table S3). The conspicuous enrichment in stress- and defense-related GO terms among the upregulated *flk-2 hos5-5* DEGs strongly suggests that *FLK* and *HOS5* cooperate to restrict the expression of stress-inducible genes. The remarkably high number of DEGs specifically upregulated in *flk-2 hos5-5* is also consistent with this view.

### Increased tolerance of *flk hos5* plants to salt and oxidative stress

In our assays on saline medium *flk-2 hos5-5* had the highest germination rate. In addition, differences in post-germination development were also observed. At 150 mM NaCl, *flk-2* and *flk-2 hos5-5* double mutants showed more cotyledons and true leaves than *hos5-5* and Col-0 (Figure S7a). However, at 200 mM NaCl, only *flk-2 hos5-5* seedlings were still able to develop open dark-green cotyledons (Figure S7b), suggesting that this background may be more tolerant to salt stress conditions. We next plated seeds on 50 mM NaCl, a concentration that did not affect germination in any of the strains examined, but impacted biomass accumulation (Figure 3a). Under these conditions, *hos5-5* seedlings appeared more affected than *flk-2* and Col-0 (Figure 3b), consistent with *hos5* sensitivity to salt stress (Chen et al., 2013). Conversely, weight-loss in *flk-2 hos5-5* plants was very moderate, suggesting that double mutants adapt better to saline/osmotic stress than single mutant and wild-type individuals (Figure 3b). These results nicely correlated with transcriptomic enrichment in *flk-2 hos5-5* of salt response genes, including, among others, *CBL-INTERACTING PROTEIN KINASE21* (*CIPK21*), *WRKY25*, *WRKY33*, or *MYB44*, whose overexpression increase Arabidopsis salt tolerance (Jung et al., 2008; Jiang and Deyholos, 2009; Pandey et al., 2015; Table S2).

**Figure 3.**
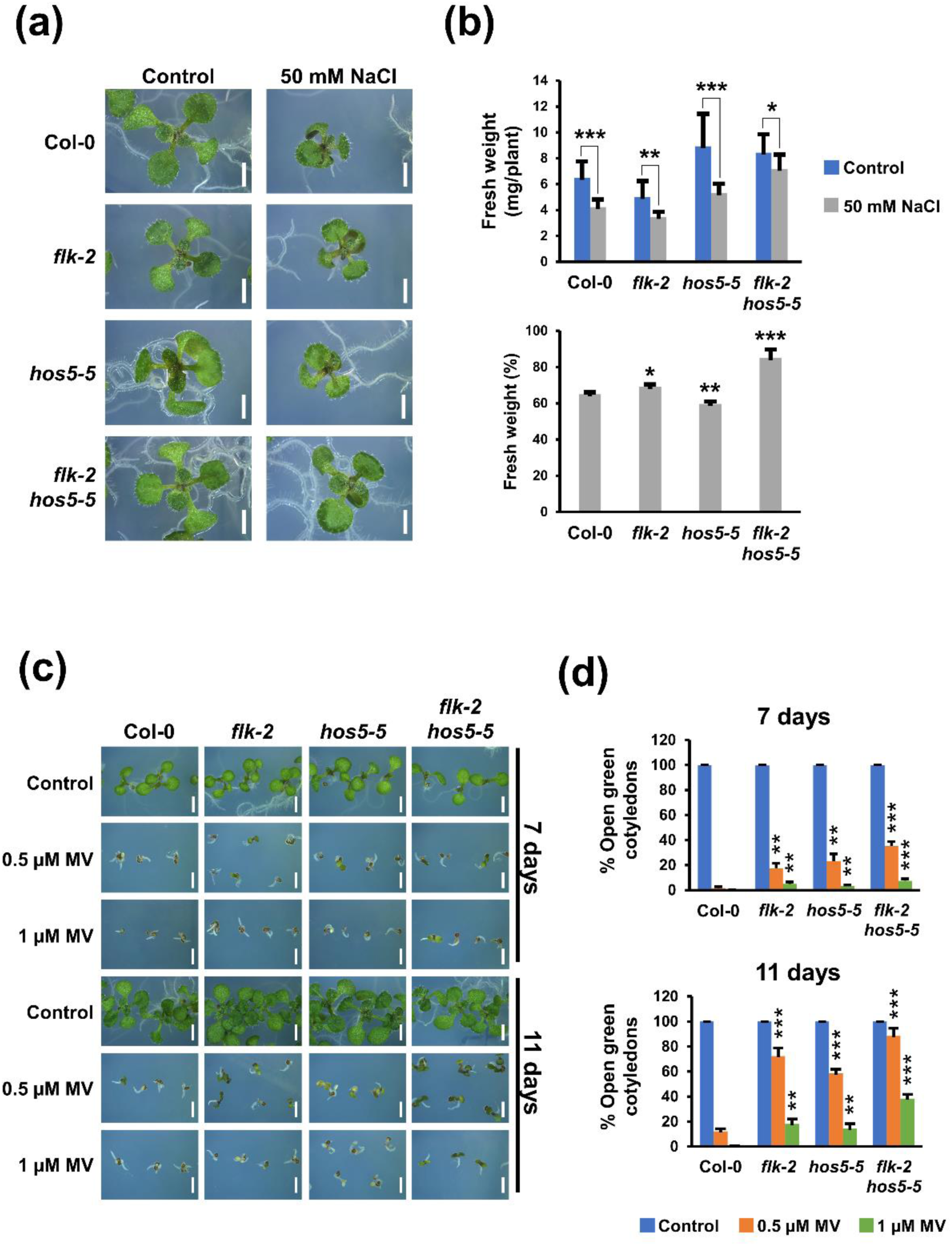
Increased tolerance of *flk hos5* plants to salt and oxidative stress. (a) Representative ten-day-old seedlings of wild-type Col-0, and mutant genotypes, grown on control medium or supplemented with 50 mM NaCl. (b) Fresh weight of ten-day-old plants grown on control medium and 50 mM NaCl. The top graph corresponds to two independent experiments with 12 plants each. Bars indicate mean ± SD and asterisks denote significant differences between control and NaCl-treated plants of the same genotype. The bottom graph represents the relative percentage of fresh weight of plants grown on NaCl with respect to their untreated controls. Asterisks indicate significant differences with respect to Col-0. (c) Wild-type Col-0 and mutant plants at 7 and 11 days after stratification, grown on control medium, or supplemented with 0.5 µM or 1 µM Methyl Viologen (MV, Paraquat), respectively. (d) Percentage of open green cotyledons in germinated seeds at 7 and 11 days in control medium, or in the presence of MV. Data correspond to three independent experiments with no less than 100 seedlings per genotype. Asterisks indicate significant differences with respect to untreated controls of the same genotype. *, *P* < 0.05; **, *P* < 0.01; ***, *P* < 0.001. Scale bars, 0.2 cm.

Detrimental effects of salt stress, aside from osmotic and ionic imbalance, often lead to reactive oxygen species (ROS) production and subsequent oxidative damage, a secondary effect common to other types of stress (Fichman & Mittler, 2020; Mansour & Hassan, 2022). Therefore, we tested sensitivity to the oxidative stress inducer methyl viologen (MV). Based on the percentage of established seedlings with green open cotyledons, all *flk*/*hos5* mutant backgrounds were less sensitive to MV when compared to Col-0 (Figure 3c,d). Indeed, resistance of *flk-2* to oxidative stress was previously reported (Fabian et al., 2023). However, although the GO term ‘response to oxidative stress’ was enriched in all three mutant backgrounds (Table S3), *flk-2 hos5-5* plants showed the most robust resistance (Figure 3c,d). This paralleled the increased expression of numerous antioxidant activities, including peroxidases, glutathione S-transferases and catalases, some of which were specifically upregulated in the double mutant (Table S2). Additional upregulated genes known to promote oxidative stress tolerance included *WRKY25*, *WRKY33*, *CIPK9*, or *PATELLIN2* (*PATL2*) (Hornbergs et al., 2023; Figure S6; Table S2). Stress-derived ROS production mostly depends on the NADPH oxidase RESPIRATORY BURST OXIDASE HOMOLOG D (RBOHD), which is activated by BOTRYTIS INDUCED KINASE 1 (BIK1) and negatively regulated by the recently described PHAGOCYTOSIS OXIDASE/BEM1P (PB1) DOMAIN-CONTAINING PROTEIN

(PB1CP) (Goto et al., 2024). Interestingly, all were found to be upregulated in the *flk-2 hos5-5* mutant, likely contributing to fine-tune the final output of ROS production (Table S2). These assays functionally validate our RNA-Seq data and further support a role for *FLK* and *HOS5* in abiotic stress responses.

### The *flk hos5* double mutant exhibits augmented SA and JA levels and resistance to fungal infection

The expression profiles of loci related to SA and JA biosynthesis/signaling pathways displayed clear differences between *flk-2* and *hos5-5* single mutants. For example, the expression of SA-related genes, such as *PHYTOALEXIN DEFICIENT 4* (*PAD4*), *ISOCHORISMATE SYNTHASE 1* (*ICS1*) and *ACCELERATED CELL DEATH 6* (*ACD6*) (Dempsey et al., 2011), decreased in *flk-2*, whereas in *hos5-5* plants increased or they remained unchanged. Accordingly, the SA marker *PATHOGENESIS-RELATED GENE 1* (*PR1*) (Jung & Hwang, 2000) was downregulated in *flk-2* but overexpressed in *hos5-5* (Table S2). These results agree with *FLK* positive regulation of SA-mediated defense (Fabian et al., 2023).

On the other hand, JA-associated GO terms were overrepresented among *hos5-5* downregulated genes (Table S3), as observed in the allelic mutant *esr1-1* (Thatcher et al., 2015). Conversely, these terms were enriched among *flk-2* upregulated activities, including key genes for JA biosynthesis or signaling such as *LIPOXYGENASE 2* (*LOX2*), *LOX3*, *ALLENE OXIDE SYNTHASE* (*AOS*), *ALLENE OXIDE CYCLASE 2* (*AOC2*), *MYC2*, and *VEGETATIVE STORAGE PROTEIN 1* (*VSP1*) (Wasternack & Hause, 2013; Table S2).

Antagonistic interactions between JA and SA are well documented in Arabidopsis (Hou & Tsuda, 2022). However, *flk-2 hos5-5* double mutants seemed, interestingly, to recapitulate features from both single mutants. Some SA key genes were upregulated (e.g. *PAD4*) or unchanged (e.g. *ICS1*), but still maintaining high levels of *PR1* expression (Table S2). Notably, the SA receptor-encoding gene *NONEXPRESSOR OF PR GENES 1* (*NPR1*) (Zavaliev & Dong, 2024) was significantly upregulated only in *flk-2 hos5-5* plants (Table S2). On the other hand, JA-related genes, including two of the most characteristic JA-activity markers, such as *MYC2* and *PDF1.2* (Wasternack and Hause, 2013), also showed high transcript abundance in *flk-2 hos5-5*, as well as key genes for plant growth-defense trade-off under JA signaling (e.g. *MYB44*, *WRKY18*, *WRKY33*, *ORA47*) (Zhang et al., 2020b; Wang & Zhang, 2021; Table S2).

SA and JA play crucial roles in plant immunity against biotrophic/hemibiotrophic and necrotrophic pathogens, respectively (Zhang et al., 2020c). Molecular signatures for SA and JA activities in our transcriptomic dataset prompted us to test the susceptibility of *flk*/*hos5* mutants to phytopathogenic fungi with different lifestyles. Plants were inoculated with the hemibiotroph *Fusarium oxysporum*, responsible for wilt disease. Col-0 and *flk-2* mutants did not show significant differences when exposed to *F. oxysporum* (Figure 4a,b), and their endogenous SA levels were also very similar (Figure 4c), despite lower expression of key SA-related genes in *flk-2*. By contrast, the *hos5-5* mutant exhibited higher endogenous SA levels than Col-0, although higher resistance to *F. oxysporum* was not statistically significant, probably due to interassay variability in this mutant (Figure 4b,c). Actually, the *hos5* allele *esr1-1* was reported to be more resistant to wilt disease (Thatcher et al., 2015). Interestingly, the *flk-2 hos5-5* double mutants were clearly more resistant to *F. oxysporum* (Fig 4b), and their endogenous SA levels were significantly higher than those of *hos5-5* plants (Fig 4c).

**Figure 4.**
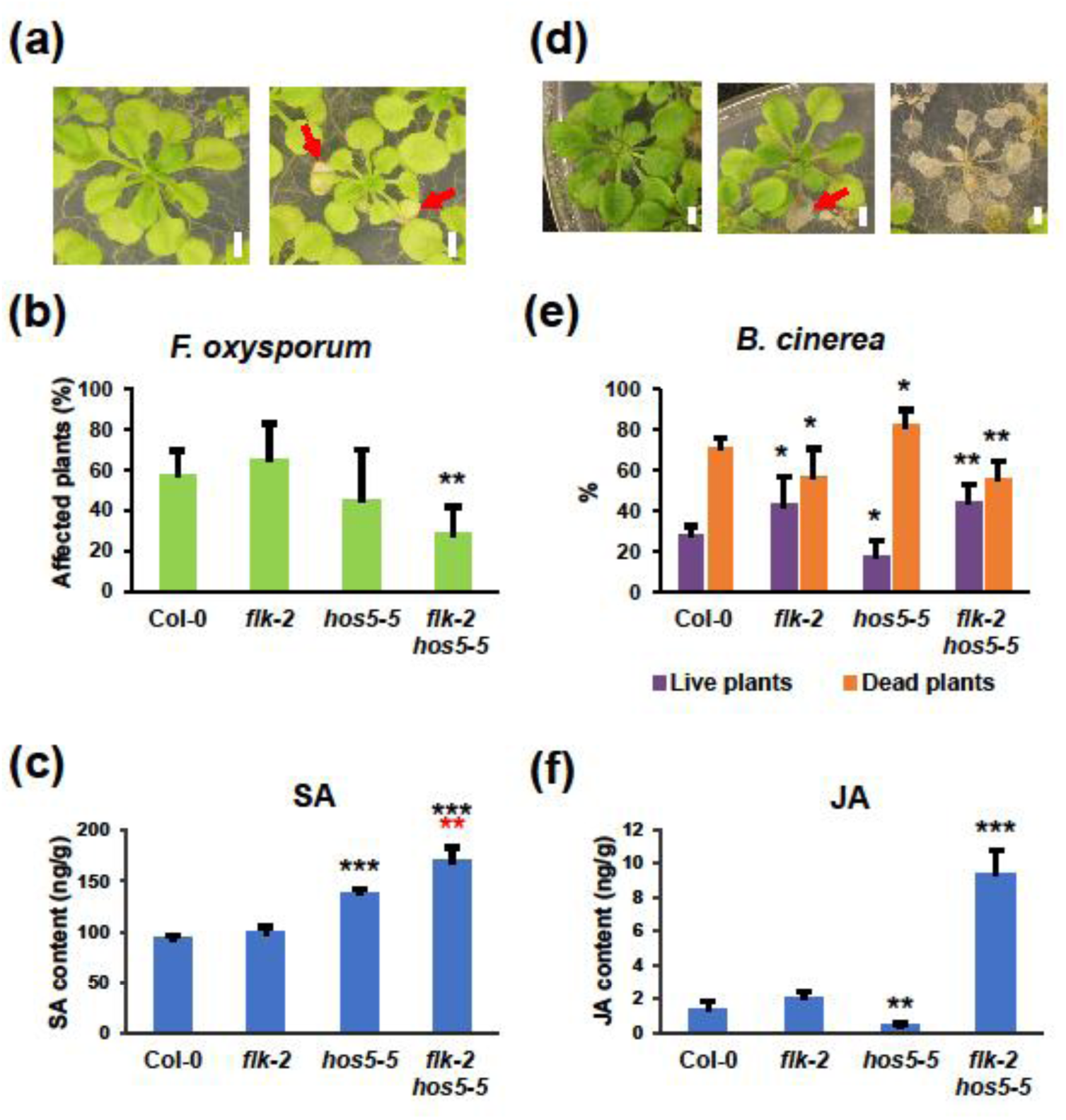
Increased resistance of the *flk hos5* mutants to fungal pathogen infection. (a) Representative asymptomatic (left) and affected (right) two-week-old plants infected with *F. oxysporum* photographed at 15 dai (days after infection). (b) Average percentage of plants showing *F. oxysporum* disease symptoms at 15 dai. (c) Total SA content in 12-day-old wild-type and mutant plants. (d) Representative two-week-old plants infected with *B. cinerea* at 15 dai. Asymptomatic plants (left), live plants with necrotic lesions (middle), and dead plants (right). (e) Average percentage of live (purple) and dead (orange) plants infected with *B. cinerea* at 15 dai. (f) Total JA content in 12-day-old wild-type and mutant plants. For fungal inoculations, three biological replicates were carried out, each containing at least forty plants per genotype. For hormone measurement, data represent the mean value of three replicates with at least 30 plants per sample. Bars indicate mean ± SD. Black and red asterisks indicate significant differences with respect to corresponding Col-0 controls and *hos5-5* plants, respectively (*, *P* < 0,05; **, *P* < 0,01; ***, *P* < 0.001). Red arrows in (a) and (d) indicate disease lesions in live plants. Scale bars, 2mm.

We also challenged our mutant strains with *Botrytis cinerea*, a necrotrophic fungus causing gray mold disease in many plant species, including crops (Bi et al., 2023). The ratio between dead and live plants indicated that *hos5-5* was more susceptible than Col-0, whereas *flk-2* and *flk-2 hos5-5* mutants were more resistant (Figure 4d,e). Increased resistance to *B. cinerea* by *flk* nicely fits with the enrichment of JA-related GO terms in *flk-2* and *flk-2 hos5-5* (Table S3), and it has been recently reported by another group (Fabian et al., 2023). Also consistent with transcriptomic data and disease severity, JA levels in *hos5-5* were significantly lower than those of Col-0 (Figure 4f). Remarkably, the endogenous JA level in *flk-2 hos5-5* double mutants was very high, exceeding by far that of *flk-2* plants (Figure 4f). This was surprising because, despite such a difference in JA content, resistance to *B. cinerea* was very similar in both mutants (Figure 4e,f).

High investment in defense is usually associated with growth or developmental alterations (Karasov et al., 2017; Zhang et al., 2020a). However, no morphological anomalies were detected in *flk-2 hos5-5*, prompting us to consider uncoupling of stress and growth limitation. We therefore decided to monitor primary root growth. Plants with high endogenous JA levels typically exhibit a short-root phenotype and sensitivity to the JA-derivative methyl-jasmonate (MeJA) (Wasternack & Hause, 2013). However, *flk-2 hos5-5* roots were similar to those of Col-0 (Figure S8a). Additionally, mutant and wild-type roots did not differ much when exposed to increasing MeJA concentrations. Only at 50µM MeJA, *flk-2 hos5-5* roots were slightly shorter (Figure S8). These results suggest that stress tolerance and growth restriction might be uncoupled, as previously postulated for *hos5* (Thatcher et al., 2015). The growth inhibitory effect of JA might be counteracted by other misregulated gene activities. For instance, the JA negative regulator *JAM1*/*bHLH17*, whose overexpression attenuates JA-mediated root inhibition (Han et al., 2023), is upregulated in the three mutant backgrounds (Table S1). Likewise, endogenous JA levels might be modulated by negative feedback mechanisms (Gasperini & Howe, 2024). Accordingly, some genes encoding JA catabolic enzymes, such as *SULFOTRANSFERASE 2A* (*ST2A*), which is enhanced by JA treatments (Gidda et al., 2003), were also upregulated in *flk-2 hos5-5* (Table S2).

### mRNA regulation mediated by *FLK* and *HOS5*

Our results corroborate that *FLK* and *HOS5* act in concert to control floral transition through *FLC* regulation, and strongly suggest that they also orchestrate stress-responses by regulating additional genes, including stress-related loci (see above). To gather information on how *flk* and *hos5* mutations jointly impact mRNA expression, we first explored *FLC* regulation*. FLK* has been reported to affect *FLC* splicing efficiency (Amara et al., 2023). We therefore monitored spliced and unspliced *FLC* transcripts corresponding to the large intron 1, common to all *FLC* isoforms, and the terminal intron 6 of *FLC* variant 1 (Figure 5a). In *flk-2*, the levels of spliced products increased significantly more than their respective unspliced forms (Figure 5b,c), being splicing efficiency (ratio of spliced over unspliced transcripts) higher than in Col-0 (Figure 5d). This aligns with recent findings indicating that FLK binds to *FLC* mRNA in an m6A-dependent manner to impair splicing (Amara et al., 2023). On the other hand, levels of correctly spliced forms in the *hos5-5* mutants were slightly lower than those in Col-0 which, together with a modest increment of unspliced forms, led to reduced splicing efficiency as a result (Figure 5b-d). In fact, splicing efficiency of both introns in *flk-2 hos5-5*, although higher than in Col-0, was lower than that of *flk-2*, despite the notable increase of spliced products (Figure 5b-d). *HOS5* affects splicing but also impairs transcription by preventing 5’ capping (Chen et al., 2013; Jiang et al., 2013). Therefore, and as previously reported for *flk*, high *FLC* abundance may result from increased splicing efficiency (Amara et al., 2023). Nevertheless, enhanced transcription and/or increased transcript stability cannot be ruled out in *flk-2 hos5-5* plants.

**Figure 5.**
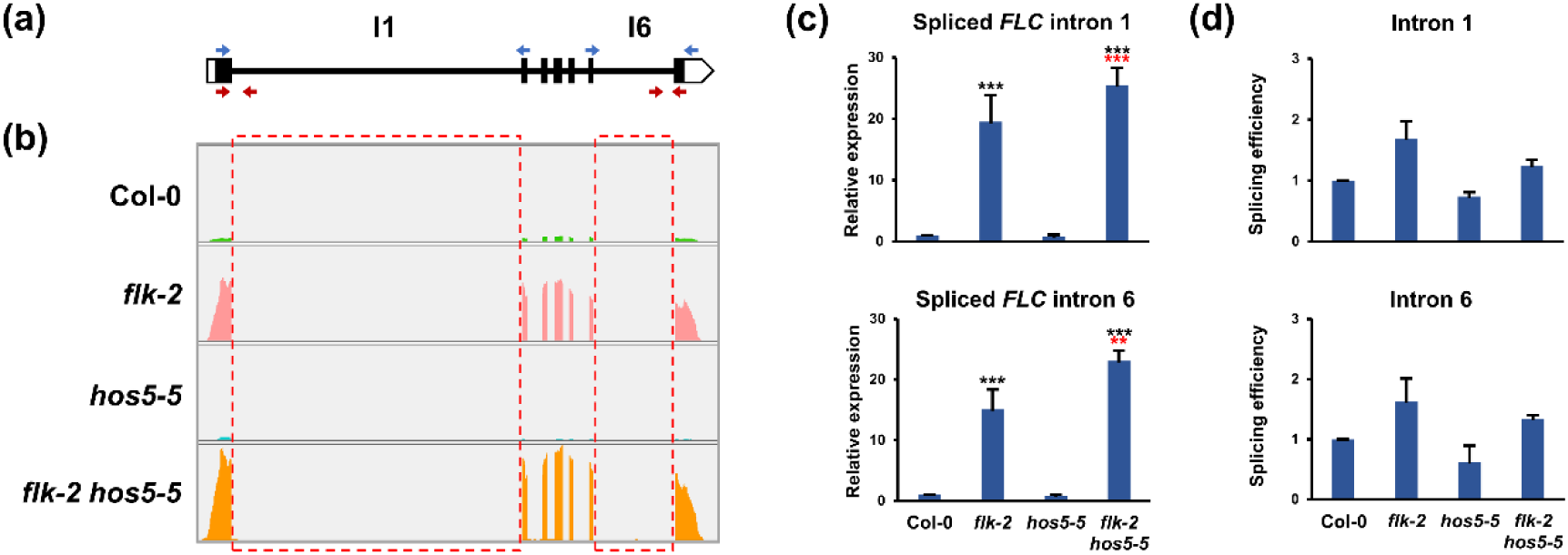
Splicing efficiency of *FLC* pre-mRNA in the *flk-2*, *hos5-5* and *flk-2 hos5-5* mutants. (a) Schematic representation of the *FLC* gene (isoform 1). Thick bars indicate exons (black, translated; white untranslated). Thin lines denote introns. Blue and red arrowheads indicate positions of primers used to amplify spliced and unspliced products, respectively. The numbers of examined introns are indicated on top. (b) Wiggle plots of *FLC* RNA-Seq data in Col-0 and indicated mutant backgrounds. Read counts were normalized as determined by IGV software. The dashed boxes indicate introns analyzed in panels (c) and (d). c) Relative expression of *FLC* monitored by qRT-PCR as the amount of spliced intron 1 (top) or spliced intron 6 (bottom) transcripts. (d) Splicing efficiency (measured as the ratio of the accumulation of spliced forms to unspliced forms) of introns 1 and 6, respectively. For (c) and (d) panels, data correspond to three biological replicates with three technical replicates each. Bars represent mean ± SD. Black and red asterisks indicate significant differences with respect to Col-0 and *flk-2* plants, respectively (**, *P* < 0.01; ***, *P* < 0.001).

To gain a broader perspective, we further searched our RNA-seq datasets for introns differentially expressed in our mutants relative to the wild type. In *flk-2*, we found 228 differentially upregulated introns (intron-specific reads more abundant than in Col-0), corresponding to 97 genes, 37 of which were upregulated in this mutant, whereas only one was downregulated (Figure 6; Figure S9a; Figure S10; Table S4). Likewise, 119 introns, representing 47 genes, were upregulated in *hos5-5*. Among the genes involved, 33 were upregulated in *hos5-5*, and none was downregulated (Figure 6; Figure S9a; Figure S10; Table S4). Remarkably, we found 1328 upregulated introns in the *flk-2 hos5-5* double mutant, significantly more than in either single mutant. This set represented 450 genes, 301 of which were upregulated in *flk-2 hos5-5*, being most of them double-mutant specific (253, 84%) (Figure 6; Figure S9a; Figure S10; Table S4). By contrast, we detected only 9 downregulated genes in this group, including that found among *flk-2* downregulated genes (Figure S9a).

**Figure 6.**
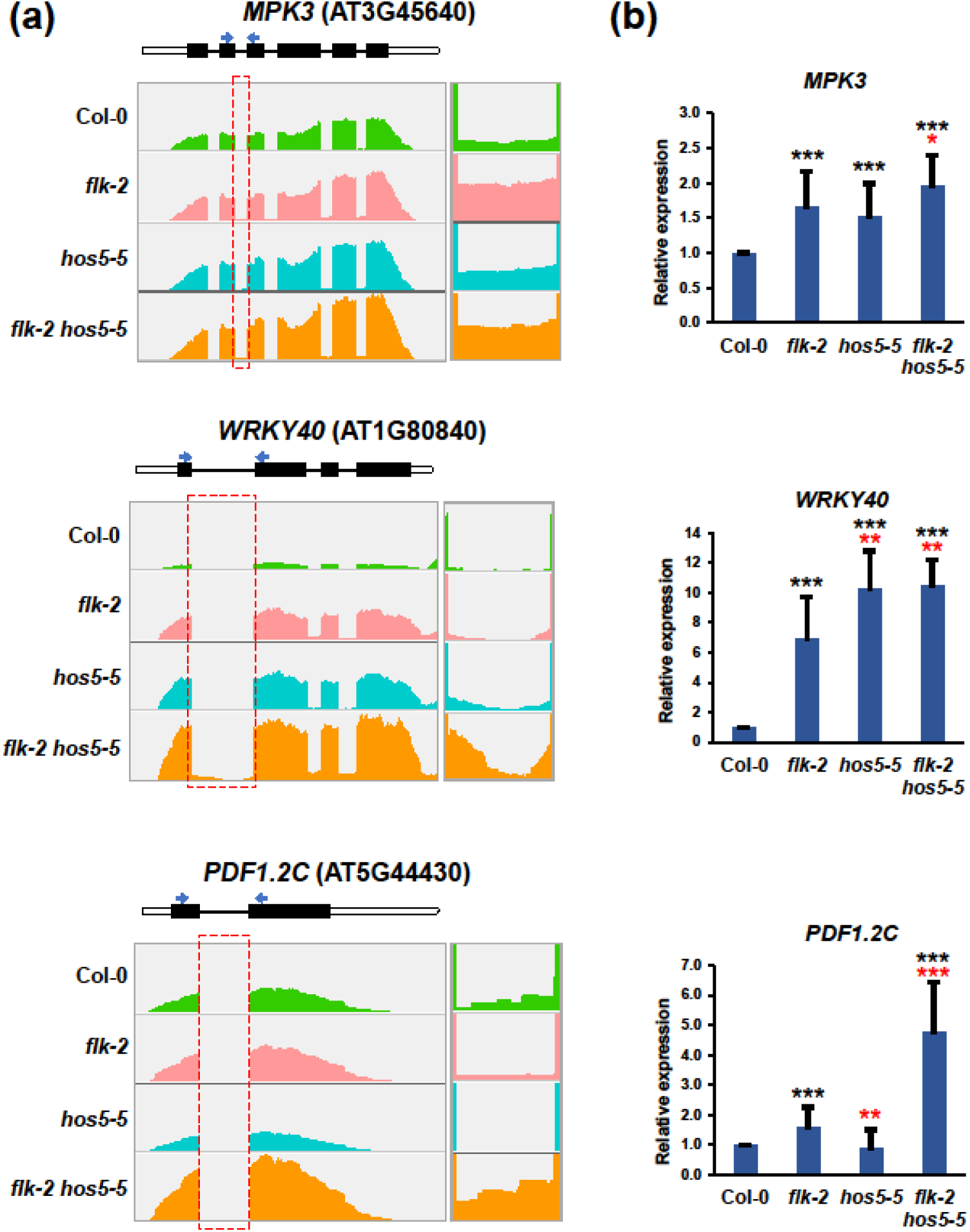
Upregulated intron sequences in upregulated genes. (a) Top of each panel: schematic representation of the corresponding gene. Thick bars indicate exons (black, translated; white untranslated). Thin lines denote introns. Arrowheads indicate positions of primers used to amplify the corresponding spliced PCR products. Bottom: wiggle plots of RNA-Seq data in Col-0 and mutant backgrounds. Read counts were normalized as determined by IGV software. The dashed boxes indicate introns analyzed in the right panels, a magnification of which is shown on the right. In these examples, indicated intron-specific reads parallel the mRNA expression level in each genotype. (b) Relative gene expression monitored by qRT-PCR as the amount of indicated spliced transcripts. Data correspond to three biological replicates with three technical replicates each. Bars represent mean ± SD. Black and red asterisks indicate significant differences with respect to Col-0 and *flk-2* plants, respectively (*, *P* < 0.05; **, *P* < 0.01; ***, *P* < 0.001).

The above data suggest that intron retention, a frequent outcome of splicing alteration in plants (Reddy et al., 2012), is not a prominent mechanism of gene repression in our mutant backgrounds. This agrees with previous results indicating that no significant intron retention takes place in the *hos5* mutant under normal (non-stress) conditions (Chen et al., 2013). In line with this notion, no upregulated genes harboring downregulated introns (less intron-specific reads than in Col-0) were found in any of the three mutant genotypes (Figure S9b; Figure S11; Table S4). These data indicate a positive correlation between DEGs and their differentially expressed introns. With very few exceptions, upregulated introns appeared in upregulated DEGs whereas all downregulated introns were found in downregulated DEGs (Figure S10). Taken together, our results may be consistent with FLK and HOS5 actions on mRNA maturation, including splicing, since higher splicing efficiency of featured introns in the *flk-2* mutant was observed in some upregulated genes (Figure S6; Figure S12), but may also reflect additional effects on transcript stability and/or transcription rate, particularly in *flk-2 hos5-5* plants.

### FLK and HOS5 physically interact

*FLK* and *HOS5* encode KH-domain polypeptides that regulate transcription and RNA processing of their target genes (Jeong et al., 2013; Jiang et al., 2013; Rodríguez-Cazorla et al., 2018; Amara et al., 2023). The results described above indicate that both proteins are functionally related. We previously identified other KH-protein partners of FLK (Rodríguez-Cazorla et al., 2015) and, therefore, sought to determine whether FLK and HOS5 could associate. Indeed, our *in planta* bimolecular fluorescence complementation (BiFC) confirmed an interaction between FLK and HOS5 in leaf cell nuclei (Figure 7a), consistent with the localization of the individual proteins (Mockler et al., 2004; Jiang et al., 2013). This interaction was further corroborated by yeast-two-hybrid (Y2H) assays (Figure 7b). Together, these findings strongly support a functional interplay between FLK and HOS5.

**Figure 7.**
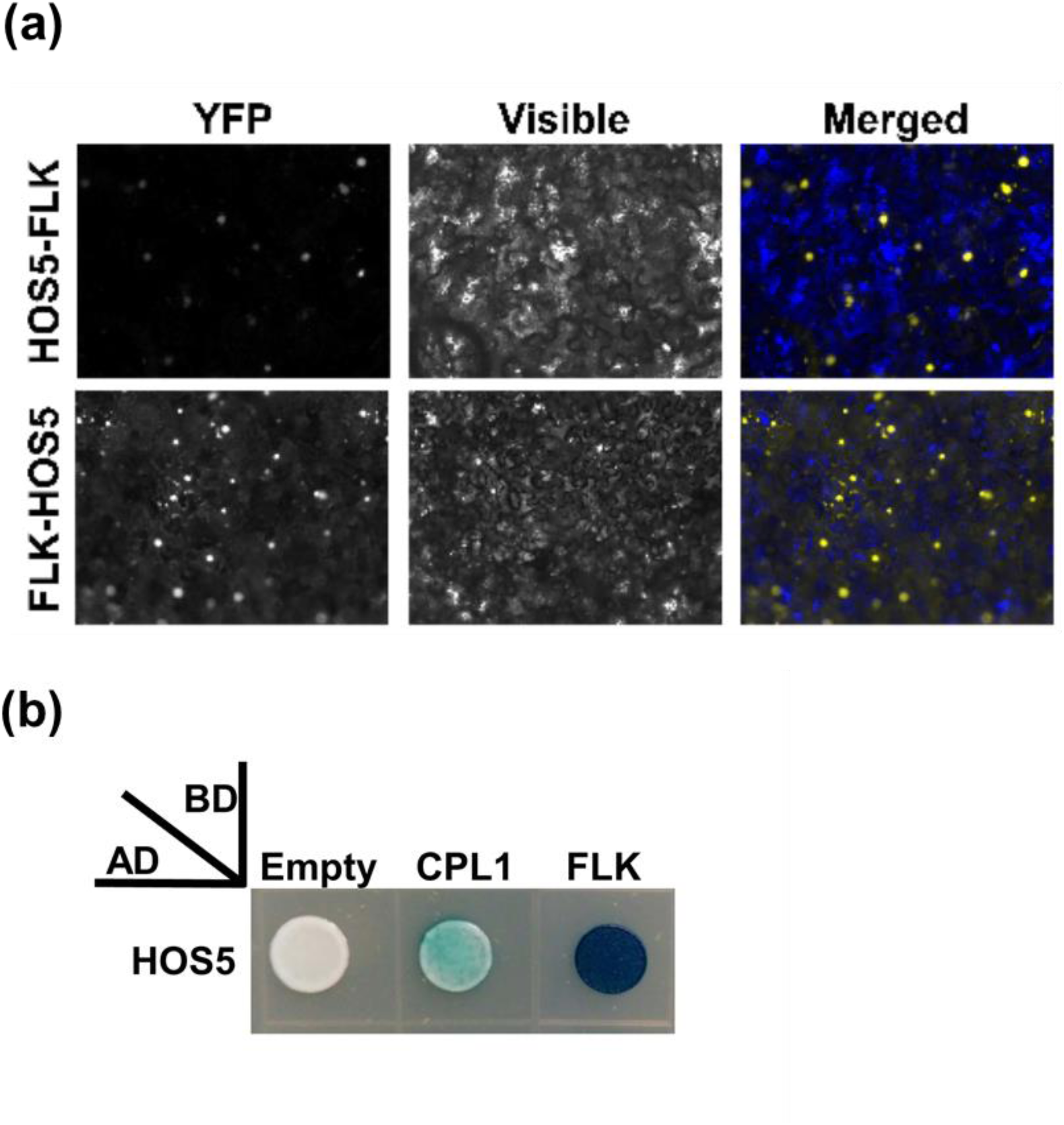
FLK and HOS5 *in vivo* interaction. (a) BiFC visualization of protein dimerization (yellow fluorescence) in *Nicotiana benthamiana* leaf cells agroinfiltrated with constructs encoding HOS5 and FLK fusion proteins. In each test, the first protein was fused to the C-terminal fragment of the YFP (YFPct), and the second protein to the N-terminal portion (YFPnt), respectively. In the merged (visible+YFP fluorescence) picture, yellow nuclei are seen on blue background used to increase contrast. (b) FLK-HOS5 interaction in yeast two-hybrid assays. CPL1-HOS5 interaction was previously reported and used as a positive control (Chen et al., 2013; Jeong et al., 2013; Jiang et al., 2013).

## Discussion

Plants have evolved the ability to adjust their flowering time to environmental fluctuations and stress conditions (Kazan & Lyons, 2016). Regulatory genes involved in floral timing and stress are crucial, likely serving as key molecular hubs to adequately integrate these responses. Here, we provide evidence based on genetic, molecular, physiological and transcriptomic analyses, that delineate the KH-domain genes *FLK* and *HOS5* as an integrating regulatory node that couples flowering and stress responses to secure plant survival and reproductive success.

### *HOS5* cooperates with *FLK* to repress *FLC* and promote flowering

We have demonstrated that *HOS5* strongly interacts with *FLK* to promote the floral transition via *FLC* repression. The role of *HOS5* as a floral regulator has been previously overlooked due to the weak effect of *hos5* mutant alleles on flowering. In contrast, *hos5* led to a synergistic increase of *FLC* levels and flowering time when combined with *flk*. Supporting its role in flowering, *HOS5* shows expression peaks in the vegetative and reproductive apices (Karlsson et al., 2015). Moreover, the contribution of *hos5* to *FLC* overexpression was also evidenced by the enhanced *FLC*-dependent germination rates of *flk-2 hos5* seeds under salt stress.

### *FLK* and *HOS5* jointly modulate the expression of stress inducible genes

Aside from *FLC*, our RNA-Seq data analysis revealed a substantial overlap of DEGs between the *flk-2* and *hos5-5* single mutants. Additionally, DEGs found in *flk-2 hos5-5* were about three times those in either single mutant, most of them specific to the double mutant. This interesting feature and the enrichment in stress-related GO terms, suggests that, besides their independent roles, co-regulation via FLK-HOS5 plays an important role in the modulation of stress-related genes. It is tempting to anticipate that a fraction of them could be identified as direct targets. However, additional work beyond the scope of this study is required to verify this extreme.

Prevalence of stress-related GO categories in *flk-2* and *hos5-5* agreed with previous reports for allelic mutations (Thatcher et al., 2015; Fabian et al., 2023). Remarkably, further enrichment of stress-related functions was observed among DEGs in *flk-2 hos5-5*, including terms related to salt and oxidative stress responses. In line with this, the *flk-2 hos5-5* mutant thrived better on saline medium, and showed less sensitivity to oxidative stress than Col-0 and single mutants. Numerous gene activities that confer salt tolerance and/or alleviate oxidative damage were specifically upregulated in *flk-2 hos5-5*, when comparing to single mutants. This could likely mitigate ROS-dependent deleterious effects and improve tolerance to salt stress.

Our results also suggest that, together, *FLK* and *HOS5* regulate responses to biotic agents and defense hormone homeostasis. The resistance of *hos5* to *F. oxysporum* and that of *flk* against *B. cinerea* agreed with previous studies (Thatcher et al., 2015; Fabian et al., 2023). Conversely, *hos5-5* was more susceptible to *B. cinerea*, which aligns with the reduced expression of JA-related genes and lower JA content (Figure 4). Remarkably, *flk-2 hos5-5* doubles additively combined characteristics of each single mutant: enhanced resistance to both fungal pathogens and higher levels of JA and SA. Both hormones usually act antagonistically. However, cooperative and synergistic effects have also been observed in diverse species, including Arabidopsis, indicating simultaneous activation of JA/SA signaling when required (Mur et al., 2006; Zhang et al., 2020c; Hou & Tsuda, 2022). Simultaneous deficiency of *FLK* and *HOS5* might mimic this scenario. In fact, synergistic effects of SA and JA on the expression of their respective markers *PR1* and *PDF1.2* are documented (Zhang et al., 2020c), and both genes are upregulated in *flk-2 hos5-5* (Table S2). Consistently, numerous genes involved in SA and JA responses were highly upregulated in the double mutant, potentially contributing to fungal resistance, including the key general stress regulators *ORA47* (Zeng et al., 2022) and *WRKY33*. The latter is a crucial gene for defense against necrotrophic fungi (Zheng et al., 2006), which also collaborates with the SA master regulator *NPR1* to mediate systemic acquired resistance (SAR) (Li et al., 2018; Wang et al., 2018). Recent analyses also suggest that some JA-responsive genes could enhance SA-mediated immunity, such as *MYB44*, which is upregulated in *flk-2* and, yet, significantly more abundant in *flk-2 hos5-5* (Zhang et al., 2020c; Zeng et al., 2022; Table S2).

### Stress tolerance and moderate fitness cost in *flk hos5* plants

Overexpression of “stress genes” and elevated SA and JA levels leads to increased tolerance of the *flk-2 hos5-5* double mutant to biotic and abiotic stress. However, no signs of growth limitation were observed in such plants. Uncoupled stress tolerance and growth was previously proposed for *hos5* mutants (Thatcher et al., 2015), and no evidence of impaired growth has been described for *flk* or when combined with other AP mutants (Veley and Michaels, 2008). In *flk-2 hos5-5* mutants, SA and JA activities might be regulated mainly at the signaling/perception level, perhaps favoring particular hormone branches that allow tolerance without yield cost. Alternatively, but not mutually exclusive, misregulated counteracting activities might contribute to modulate their effects. Consistent with this, root growth in Col-0 and mutants did not differ during MeJA inhibition assays, as already reported for *hos5* (Thatcher et al., 2015). Nevertheless, we cannot rule out the possibility of a poor performance by *flk-2 hos5-5* plants under additional stress conditions not evaluated in this study.

### mRNA expression regulatory mechanisms mediated by *FLK* and *HOS5*

In agreement with recent results (Amara et al., 2023), splicing efficiency of *FLC* introns increased in our *flk-2* mutants. Notably, *FLC* splicing efficiency in *flk-2 hos5-5* was lower than in *flk-2*, in spite of a much greater expression levels. In *hos5-5*, *FLC* splicing efficiency decreased, agreeing with HOS5 splicing regulation and its interaction with SR splicing factors (Chen et al., 2013). However, HOS5 also downregulates transcription by interfering with 5’ capping, which is required for efficient transcript elongation (Jiang et al., 2013). Interestingly, other *FLC* regulators, such as the RNA recognition motif proteins RZ-1B and RZ-1C, also promote efficient splicing and repress transcription (Wu et al., 2015). *FLC* overexpression in *flk-2 hos5-5* mutants may reflect dual effects on transcription and cotranscriptional processing (splicing). In the *flk-2* single mutant, increased *FLC* expression likely obeys to greater transcript stability and splicing efficiency. However, further activation of transcription in *flk-2 hos5-5* should result from efficient 5’-capping after loss of *HOS5* activity. A similar mode of action is seen for FRIGIDA (FRI), which upregulates *FLC* cotranscriptionally by direct physical interaction with the nuclear cap-binding complex (Geraldo et al., 2009). In budding yeast, slowing the RNA polymerase II elongation increases splicing efficiency whereas faster elongation reduces it (Aslanzadeh et al., 2018). This negative correlation might be adopted to explain why *FLC* (and some other genes under *FLK*/*HOS5* influence) further increases its expression levels in *flk-2 hos5-5* with respect to *flk-2* despite lower intron splicing efficiency (Figure 5).

It is likely that *FLK* and *HOS5* cooperate in additional RNA regulatory mechanisms. The lack of *HOS5* has been previously linked to altered polyadenylation site selection in some stress-inducible genes (Jiang et al., 2013), while FLK participates in *AG* transcript termination (Rodríguez-Cazorla et al., 2015). Our results indicate that FLK and HOS5 physically interact, consistent with a possible role of *FLK*/*HOS5* in regulating RNA 3’-end formation. This, in turn, might also impact on transcript stability, as demonstrated by the negative role of *FLK* on *FLC* (Amara et al., 2023).

On the other hand, our analysis of the RNA-Seq datasets suggests a minor role for splicing in explaining the global differential gene expression seen in the mutant backgrounds screened here, with very few exceptions. We observed a positive correlation between relative levels of DEGs and those of their differentially expressed introns. Thus, intron retention does not seem to be a prevailing mechanism to limit gene expression in *flk/hos5* backgrounds, at least under normal growth conditions. This was previously observed for *hos5*, in which intron retention was only detected when plants were grown on saline medium (Chen et al., 2013).

In summary, and considering all the available data, our study proposes that *FLK* and *HOS5* are part of a gene regulatory module for coordinating flowering (via *FLC* repression) and biotic/abiotic stress responses, which is likely recruited to adapt developmental responses (floral transition, reproduction, germination…) to unfavorable environmental conditions. Further dissection of the underlying regulatory mechanisms controlled by *FLK-HOS5* in modulating growth-defense status may provide valuable insights for translational strategies aimed at generating stress-tolerant crop varieties without developmental constraints.

## Supplementary information

**Figure S1.** *hos5* mutants used in this study.

**Figure S2.** Vegetative rosettes of mutant and wild-type plants.

**Figure S3.** Flowering time in *flk-2 hos5-5* under short-day conditions.

**Figure S4.** Transcript levels of *AP1, SOC1* and *TSF* in *flk-hos5* genetic backgrounds.

**Figure S5.** Venn diagrams for differentially expressed genes (DEG) in the mutants under analysis.

**Figure S6.** qRT-PCR validation of the transcriptomic RNA-Seq datasets.

**Figure S7.** Post-germinative development of *flk-hos5* mutant combinations under salt stress.

**Figure S8.** MeJA root growth inhibition assays.

**Figure S9.** Venn diagrams for differentially expressed introns and genes (DEG) in the mutants under analysis.

**Figure S10.** Upregulated intron sequences in *flk*, *hos5* and *flk hos5* upregulated genes. Additional examples.

**Figure S11.** Downregulated intron sequences in selected *flk*, *hos5* and *flk hos5* downregulated genes.

**Figure S12.** Splicing efficiency of selected genes in the *flk*/*hos5* mutants.

**Table S1.** Information about molecular genotyping, qRT-PCR, and oligonucleotides.

**Table S2.** Differentially expressed genes in *flk*, *hos5* and *flk hos5*.

**Table S3.** Overrepresented GO terms. Biological process.

**Table S4.** Differentially expressed introns in *flk*, *hos5* and *flk hos5*.

## Acknowledgements

We thank Dr. Pruneda-Paz (UCSD, USA) for critical reading of the manuscript, and Dr. Pérez-Amador (IBMCP-CSIC, Spain) for facilitating hormone measurements.

## Funding

This research was funded by the Spanish Ministry of Science, Innovation and Universities (MICIU; grant PID2020-117887GB-I00 to A.V. and A.M.-L.) and Generalitat Valenciana (CIDEGENT grant CIDEXG/2023/034 to J.J.R.).

## Author contributions

A.V., A.M.-L., J.J.R. and E.R.-C. designed the research; E.R.-C. carried out most of the experiments; J.J.R. generated constructs and performed protein assays; E.R.-C., A.A.-M. and E.Z.-G. performed fungal inoculation experiments; H.C. processed and analyzed transcriptomic data; A.V., A.M.-L., E.R.-C., and J.J.R. analyzed data; A.V. wrote the manuscript with contributions of all authors to the final draft.

## Data availability

All data supporting the findings of this study are available within the paper and online supplementary data.

## Conflict of interest

The authors declare no conflict of interest.

## Notes

### Competing Interest Statement

The authors have declared no competing interest.

