## Supplementary figures for "Integrating flowering and stress responses in Arabidopsis through KH-domain genes"

### The KH-domain genes *FLK* and *HOS5* integrate flowering and stress responses in *Arabidopsis thaliana*

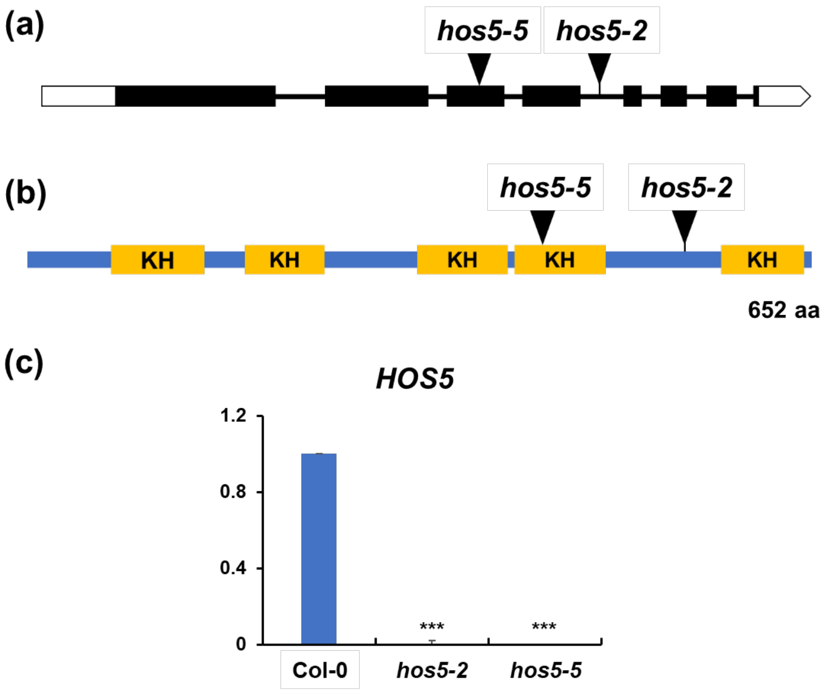


**Fig. S1.** *hos5* mutants used in this study. a) Schematic representation of the *HOS5* gene showing the location of the T-DNA insertions (triangles) corresponding to mutant alleles. Thick bars indicate exons (black, translated; white untranslated), whereas thin lines denote introns. b) The HOS5 protein, indicating its length in amino acids (aa), location of KH domains, and the relative positions of mutations with respect to the polypeptide. c) *HOS5* mRNA expression levels monitored by quantitative reverse transcription PCR (qRT-PCR) in the Col-0 wild-type and the *hos5* mutant backgrounds. Error bars: standard deviation (SD). Asterisks denote significant differences with respect to Col-0 (***, $P <$ 0.001). Data correspond to three biological replicates with three technical replicates each.

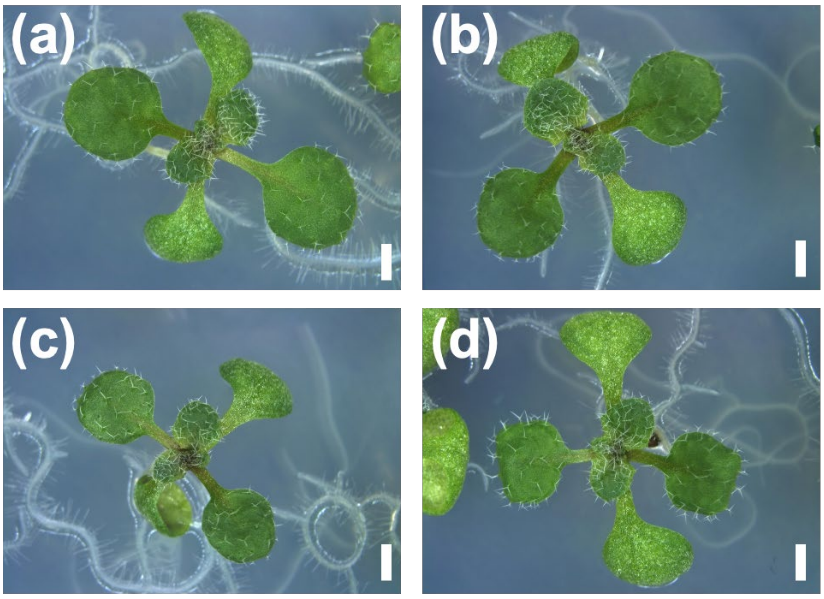


**Fig. S2.** Vegetative rosettes of mutant and wild-type plants. Nine-day seedlings of wild-type Col-0 (a), *hos5-5* (b), *flk-2* (c), and *flk-2 hos5-5* (d). Scale bar, 1 mm.

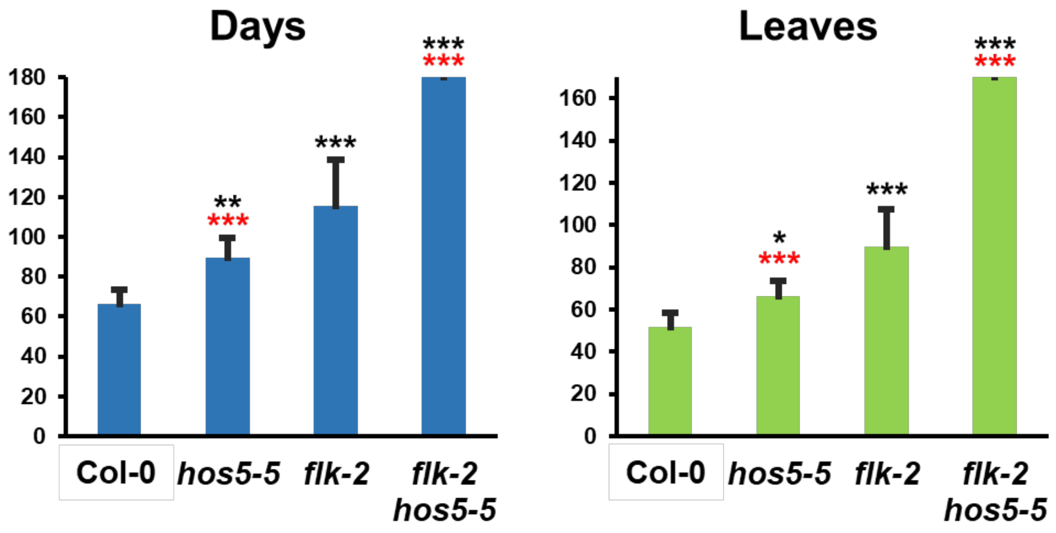


**Fig. S3.** Flowering time in *flk-2 hos5-5* under short-day conditions. Floral timing was measured as the number of days (left) or rosette leaves at bolting (right). Bars indicate mean ± SD. For each genotype 20 plants were examined. The assay was stopped at 180 days when only 4 *flk-2 hos5-5* plants had initiated flowering, thus underestimating an average of 170 leaves (bolting) for this genotype. Black and red asterisks indicate significant differences with respect to Col-0 and *flk-2* plants, respectively (*, $P < 0.05$; **, $P < 0.01$; ***, $P < 0.001$).

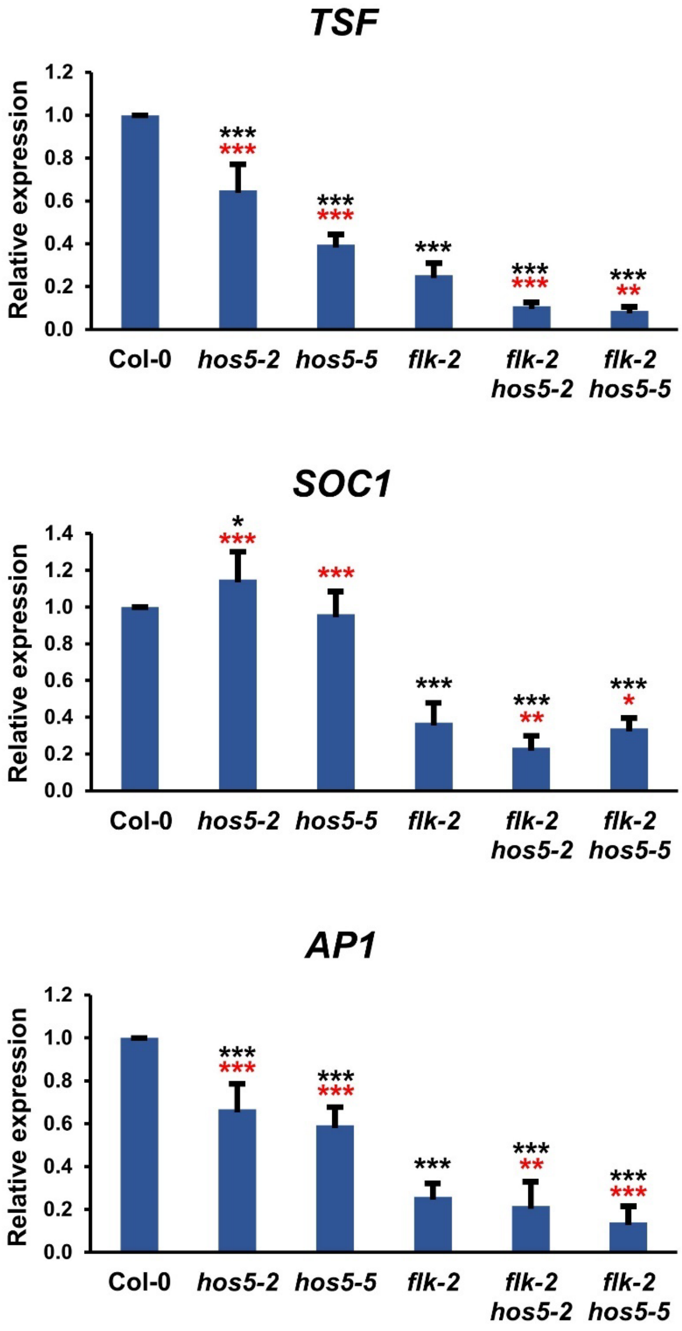


**Fig. S4.** Transcript levels of *AP1*, *SOC1* and *TSF* in *flk*-*hos5* genetic backgrounds. Relative expression, monitored by qRT-PCR, in Col-0 and corresponding mutant plants. Bars denote means ± SD. Data correspond to three biological replicates with three technical replicates each. Black and red asterisks indicate significant differences with respect to Col-0 and *flk-2* plants, respectively (*, $P < 0.05$; **, $P < 0.01$; ***, $P < 0.001$).

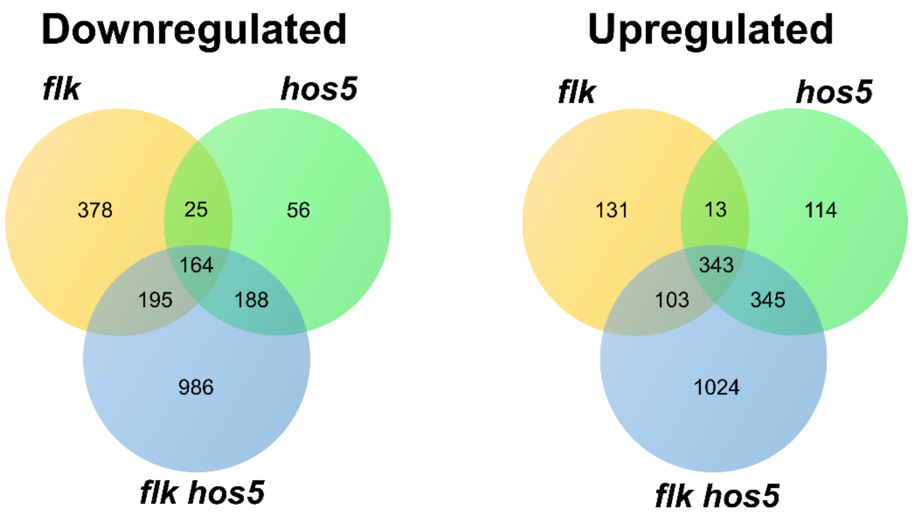


**Fig. S5.** Venn diagrams for differentially expressed genes (DEG) in the mutants under analysis. Numbers of DEGs with respect to the wild type are shown.

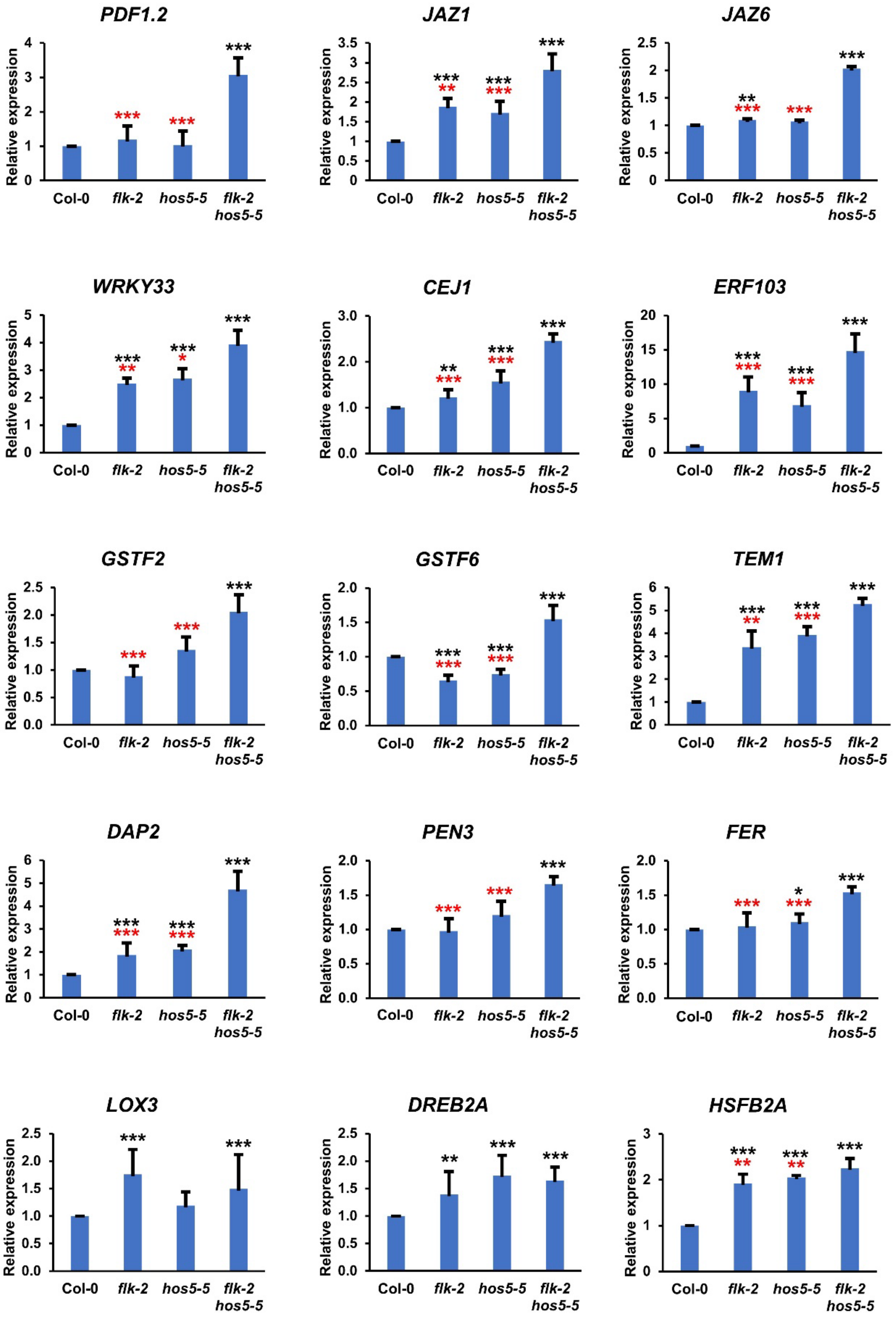


**Fig. S6**. qRT-PCR validation of the transcriptomic RNA-Seq datasets. Quantification by qRT-PCR of transcript abundance in Col-0 and mutant backgrounds of selected genes. Error bars denote SD. Black and red asterisks indicate statistically significant differences with respect to Col-0 and *flk-2 hos5-5* plants, respectively (**P* < 0.05, ***P* < 0.01, ****P* <

0.001). JA responsive genes *PLANT DEFENSIN 1.2* (*PDF1.2*) and *JASMONATE-ZIM-DOMAIN PROTEIN 1* (*JAZ1*) and *JAZ6* (Wasternack and Hause, 2013); *WRKY33*, Involved in response to various biotic (Zheng *et al.*, 2006) and abiotic stresses (Jiang and Deyholos, 2009); *COOPERATIVELY REGULATED BY ETHYLENE AND JASMONATE 1* (*CEJ1*), a DREB (subfamily A-5) ERF/AP2 transcription factor involved in defense and abiotic stress (Tsutsui *et al.*, 2009; Maruyama *et al.*, 2013); *ETHYLENE RESPONSIVE ELEMENT BINDING FACTOR 103* (*ERF103*), involved in responses to diverse stressors, including reactive oxygen species (Sewelam *et al.*, 2013; Vermeirssen *et al.*, 2014); *GLUTATHIONE S-TRANSFERASE PHI 2* (*GSTF2*) and *GSTF6*, induced by oxidative stress (Lieberherr *et al.*, 2003; Lee *et al.*, 2014); *TEMPRANILLO 1* (*TEM1*), a flowering repressor, also involved in salt responses (Osnato *et al.*, 2012); *DORMANCY/AUXIN ASSOCIATED FAMILY PROTEIN 2*/*DORMANCY ASSOCIATED GENE 2* (*DAP2*/*DRM2*), involved in negative regulation of basal defense against pathogens, SAR and ROS accumulation (Roy *et al.*, 2020); *PENETRATION 3* (*PEN3*), a plasma membrane ATP binding cassette transporter that participates in resistance to pathogens, also contributing to heavy metal resistance (He et al., 2019; Wu et al., 2019); *FERONIA* (*FER*), a multifunctional receptor-like kinase involved in ROS scavenging, ABA signaling, salt tolerance; powdery mildew infection and flowering (Wang *et al.*, 2020; Shin *et al.*, 2021; Jiang *et al.*, 2024); the JA biosynthetic gene *LOX3*; *DEHYDRATION-RESPONSIVE ELEMENT BINDING PROTEIN 2* (*DREB2A*), involved in drought stress tolerance (Sakuma *et al.*, 2006), and the *HEAT SHOCK TRANSCRIPTION FACTOR B2A* (*HSFB2A*; Wunderlich *et al.*, 2014). Underlined citations can be found in Supplementary references in Table S1.

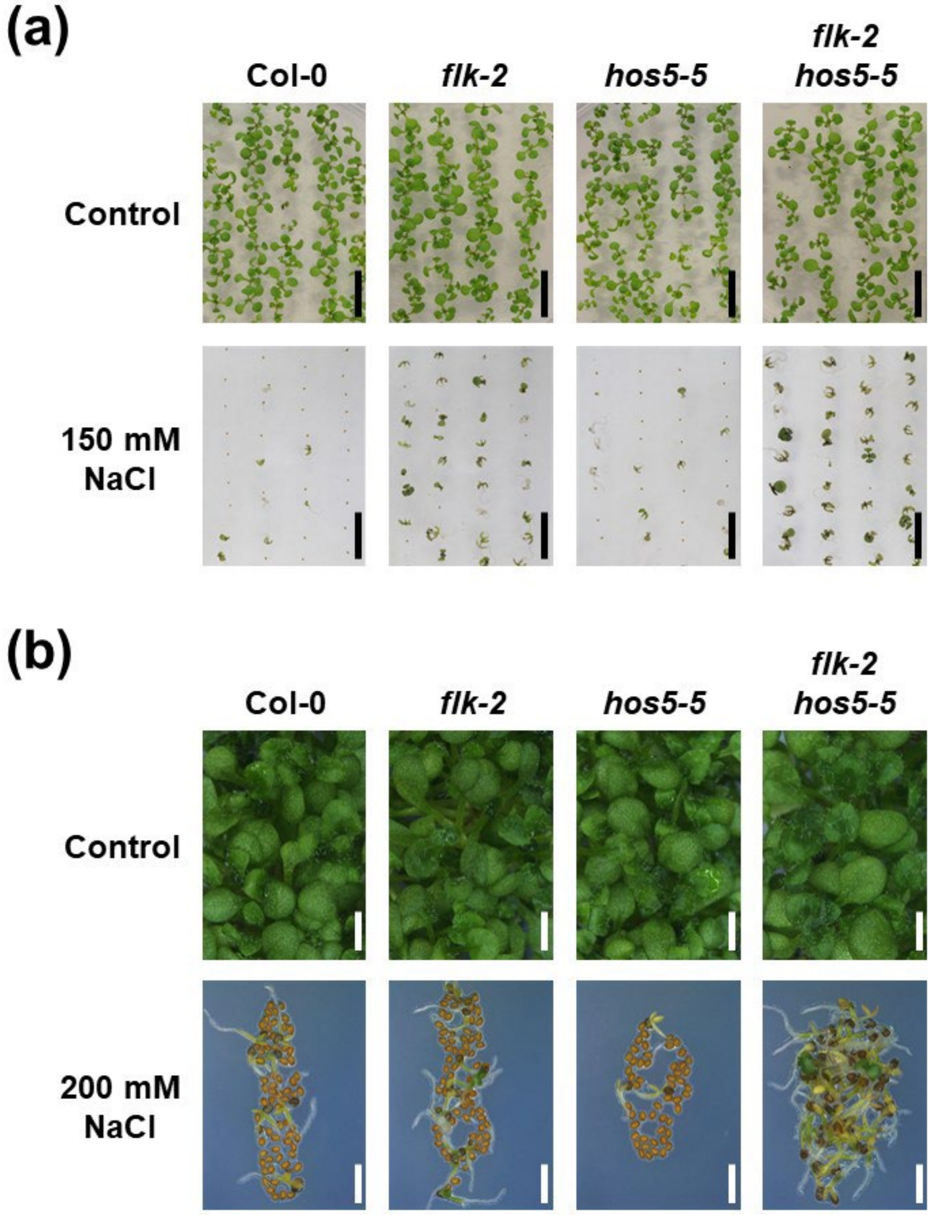


**Fig. S7.** Post-germinative development of *flk*-*hos5* mutant combinations under salt stress. (a) Thirteen-day-old seedlings grown on control medium or supplemented with 150 mM NaCl. Scale bars, 1 cm. (b) Thirteen-day-old seedlings grown on control medium or supplemented with 200 mM NaCl. Observe open green cotyledons among germinated *flk-2 hos5-5* plants. Although much smaller in size, occasional green cotyledons also developed in *flk-2* plants. Scale bars: 1 cm (a), 0.2 cm (b).

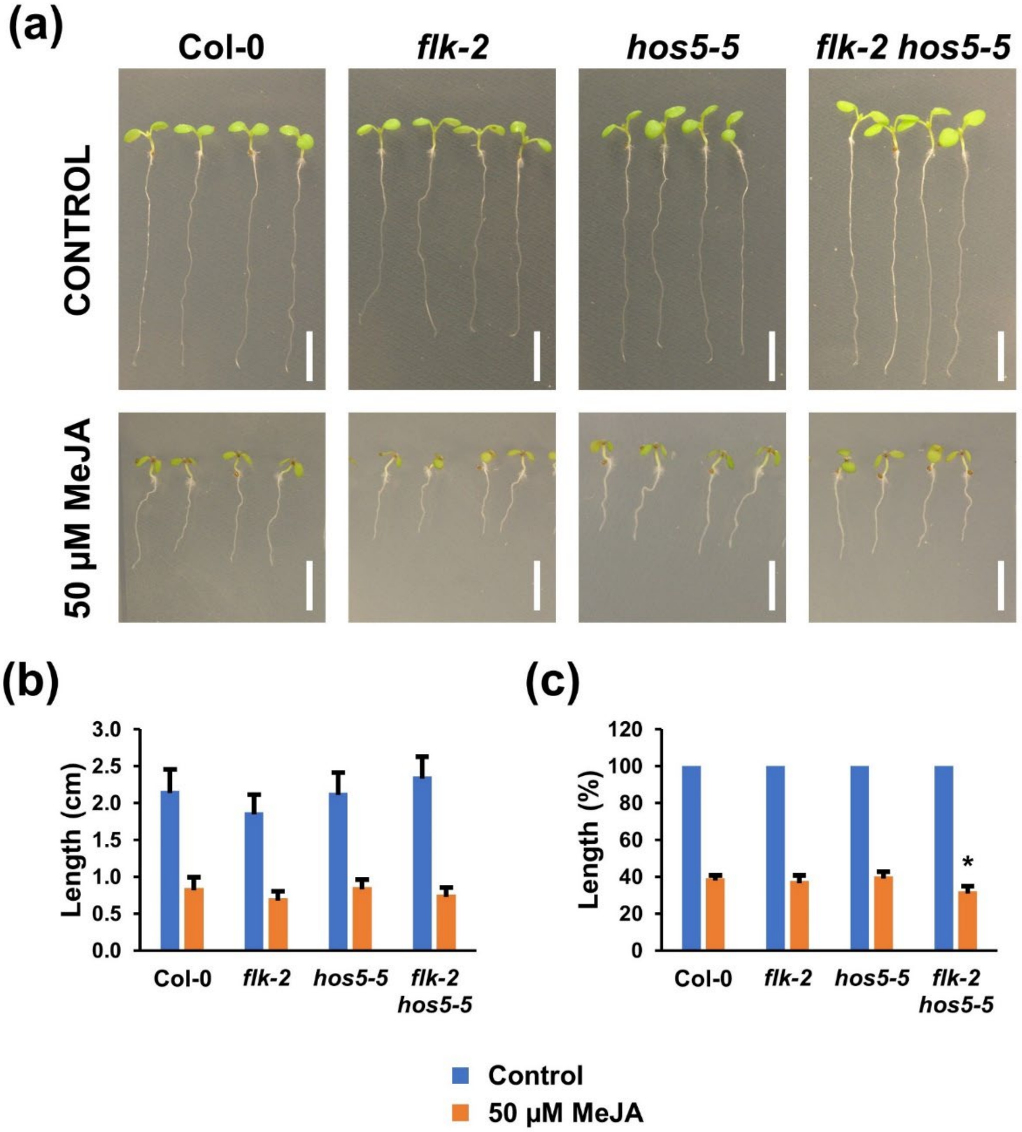


**Fig. S8.** MeJA root growth inhibition assays. (a) Seven-day-old plants grown on vertically oriented control plates (top) or supplemented with 50 µM MeJA (bottom). (b) Average length of primary root in control and MeJA-treated plants. Error bars denote ± SD of three averaged independent experiments with a minimum of 20 plants per genotype each. (c) Percentage of root length in MeJA-treated plants relative to their respective untreated controls. The asterisk in *flk-2 hos5-5* denotes a significant difference with respect to Col-0 (*, *P* < 0,05). Scale bar, 5 mm.

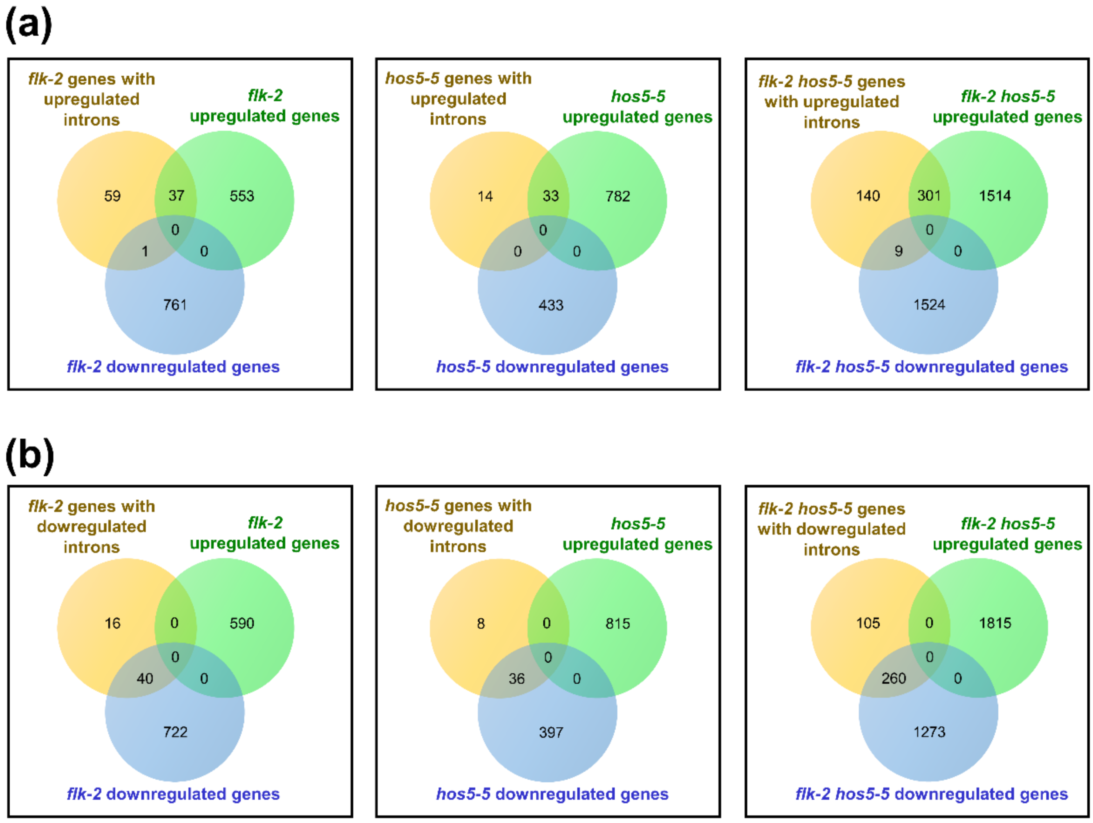


**Fig. S9**. Venn diagrams for differentially expressed introns and genes (DEG) in the mutants under analysis.
(a) Upregulated introns.
(b) Downregulated introns.
Upregulated and downregulated genes for each mutant genotype as in Table S2.

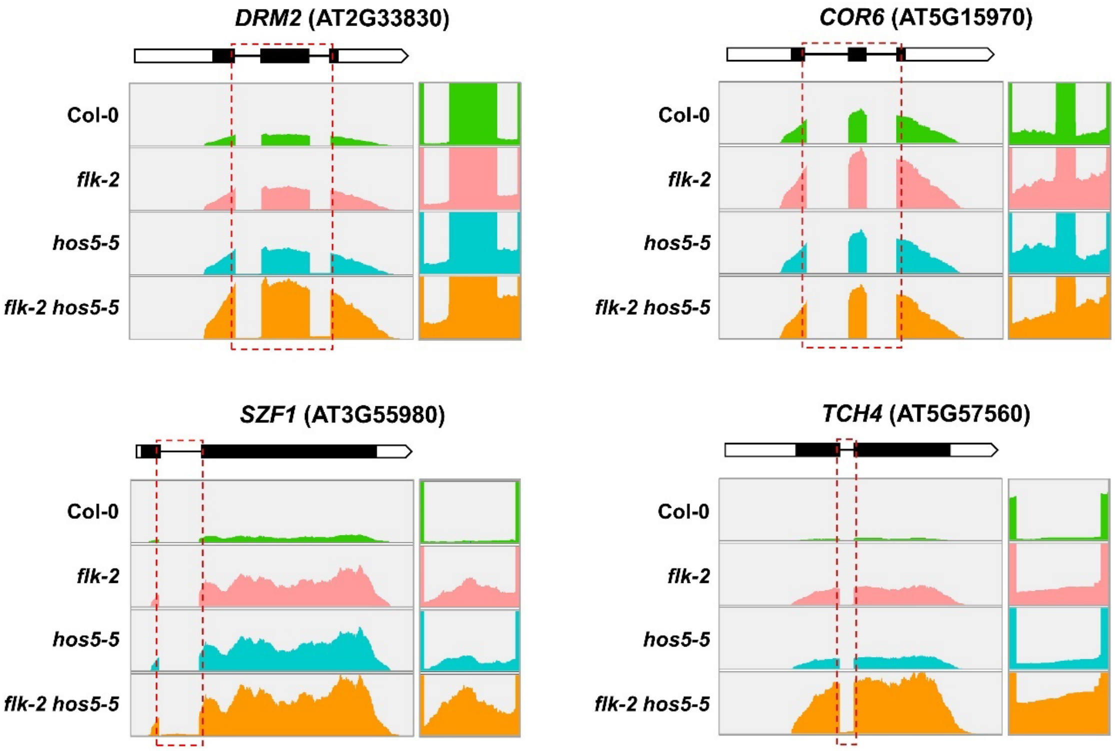


**Fig. S10.** Upregulated intron sequences in *flk*, *hos5* and *flk hos5* upregulated genes. Additional examples. Top of each panel: annotated gene structure of the corresponding gene. Thick bars indicate exons (black, translated; white untranslated). Thin lines denote introns. Bottom: wiggle plots of RNA-Seq data in Col-0 and mutant backgrounds. Read coverage is represented according to the IGV software. The red dashed boxes indicate upregulated introns, a magnification of which is shown on the right. Indicated intron-specific reads increase in parallel with the relative abundance of mRNA expression in each genotype. For *DRM2* and *COR6*, two introns are included in the same frame.

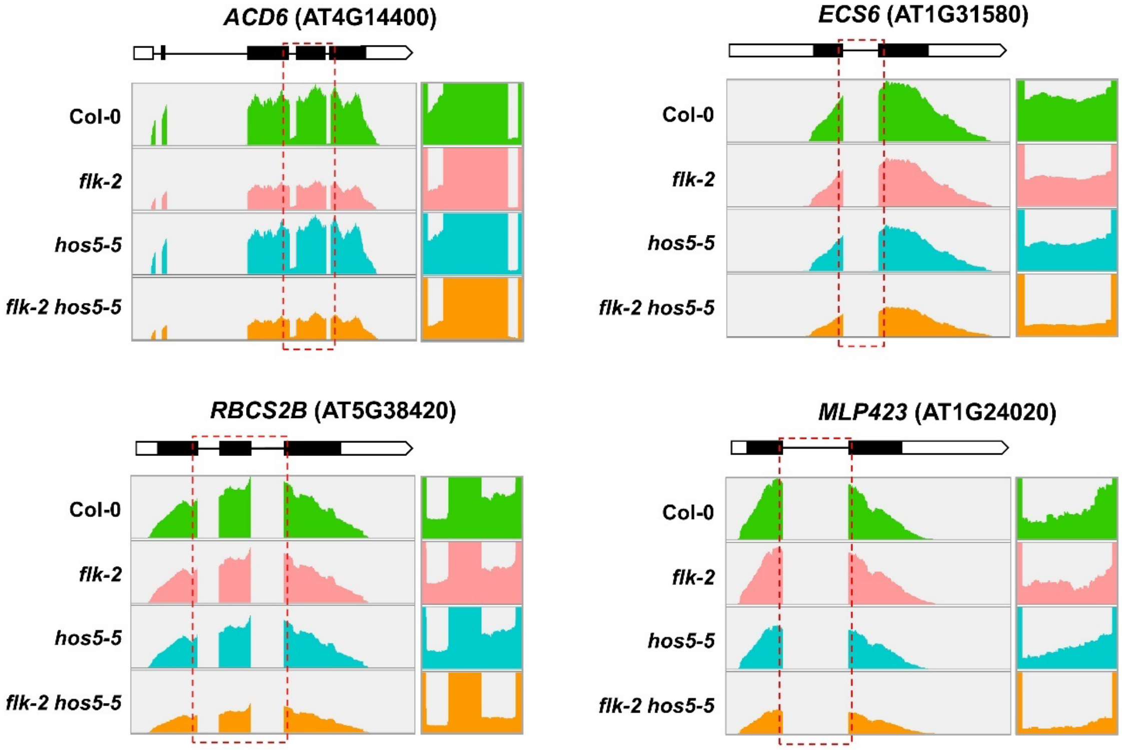


**Fig. S11.** Downregulated intron sequences in selected *flk*, *hos5* and *flk hos5* downregulated genes. Top of each panel: annotated gene structure of the corresponding gene. Thick bars indicate exons (black, translated; white untranslated). Thin lines denote introns. Bottom: wiggle plots of RNA-Seq data in Col-0 and mutant backgrounds. Read coverage is represented according to the IGV software. Featured intron areas are demarcated by red dashed frames, a magnification of which is shown on the right. In these examples, indicated intron-specific reads are less abundant in genes downregulated with respect to the wild-type. For *ACD6* and *RBCS2B* genes, two introns are included in the same frame.

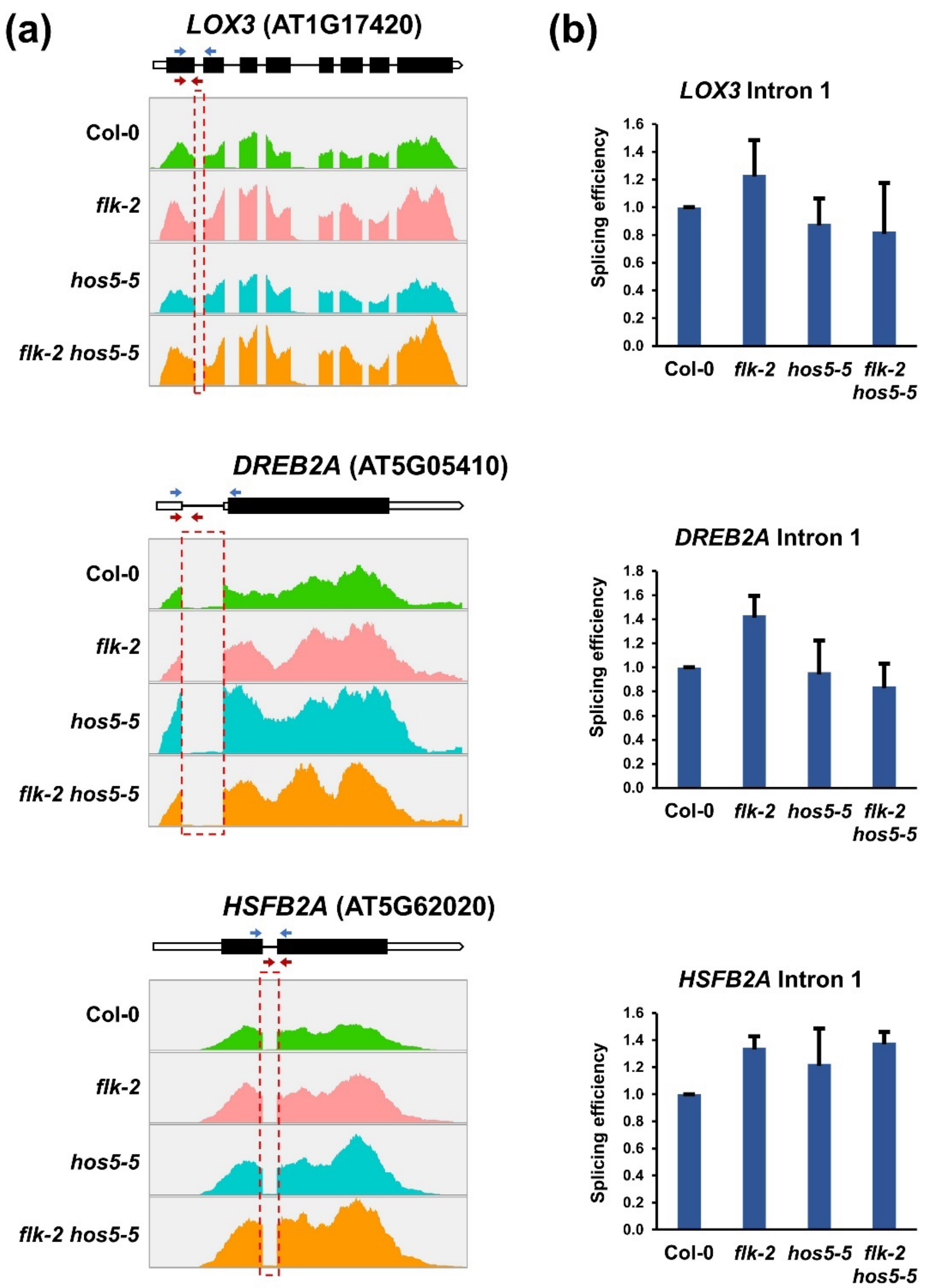


**Fig. S12.** Splicing efficiency of selected genes in the *flk*/*hos5* mutants. (a) On top of each panel: annotated gene structure of the corresponding gene. Thick bars indicate exons (black, translated; white untranslated). Thin lines denote introns. Black and red arrows indicate positions of primers used to amplify the corresponding spliced and unspliced products, respectively. At bottom: wiggle plots of RNA-Seq data in Col-0 and mutant backgrounds. Read counts were normalized as determined by IGV software. The red dashed boxes indicate introns analyzed in the right panels. (b) Splicing efficiency (measured as the ratio of the accumulation of spliced to unspliced forms). qRT-PCR relative expression data (fully spliced transcripts) are presented in Figure S6. Bars represent mean ± SD.
